# Discovery and Targeting of a Cryptic Human Proteome

**DOI:** 10.64898/2026.08.05.743124

**Authors:** Joel M. Chick, Ashley R. Woodfin, Jessica Weir, Marco Blanchette, Ariel S. Schwartz, Leopold Garnar-Wortzel, Christopher A. Polera, Sebastian C.J. Steiniger, Marcus Kelly, Kerry Wilson, Jon A. Bass, Alex M. Jaeger, Imad Ajjawi, Corey M. Dambacher

**Affiliations:** RyboDyn, Inc., Lilly Gateway Labs, 10935 Alexandria Way, San Diego, California 92121; Department of Molecular Oncology, Moffitt Cancer Center, 12902 Magnolia Dr, Tampa, Florida, 33612

**Author notes:** Contributed equally (Joel M. Chick, Ashley R. Woodfin).

## Abstract

First-in-class therapeutics require first-in-class biology. Yet despite decades of genomic and proteomic cataloging, vast regions of the human transcriptome remain dark and their encoded proteins invisible. Here we present RyboCypher™, an integrated RNA-sequencing and AI-assisted proteogenomics platform that systematically maps the RyboCypher-derived “dark” transcriptome to unannotated peptides, predicting and empirically identifying cryptic proteins across the uncharted genome. Applied to cancer cell lines, patient tumors, and matched healthy tissues, RyboCypher resolved ∼8.3 million dark RNA isoforms and ∼16 million candidate ORFs. Interrogating these against ∼0.5 billion MS/MS spectra from cellular proteomics, membrane proteomics, and immunopeptidomics datasets (comprising a total of >8,000 raw MS data files (∼7TB of MS data), derived from 2,229 patient samples), we empirically identified ∼80,000 cryptic peptides (∼10,000 cancer-associated or cancer-upregulated) at <1% FDR. Altogether, these datasets establish the CypherAtlas™, a comprehensive proteogenomic atlas of an unreported proteome comprising thousands of novel proteins, including membrane proteins with targetable extracellular domains, and intracellular proteins accessible through antigen presentation. By linking dark-RNA transcripts, predicted proteins, and patient-level metadata across RyboDyn’s proprietary experimental data, CypherAtlas further provides the training substrate for multi-modal models such as DarkCypher™, which is being developed to prioritize cryptic targets and to forecast their expression in new patient samples. As proof of therapeutic potential, we disclose evidence for a cancer-associated, cryptic protein expressed from the YBX1 locus, (cryptic YBX1; cYBX1) and demonstrate selective *in vitro* tumor cell killing through a cryptic peptide-MHC (pMHC) complex derived from this protein with a TCR-mimic (TCRm) antibody when formatted as antibody drug conjugates (ADCs). Together, RyboCypher and CypherAtlas establish the dark proteome as a vast and previously inaccessible reservoir of novel targetable biology, laying the foundation for the next generation of first-in-class therapeutics.

**Graphical Abstract:** 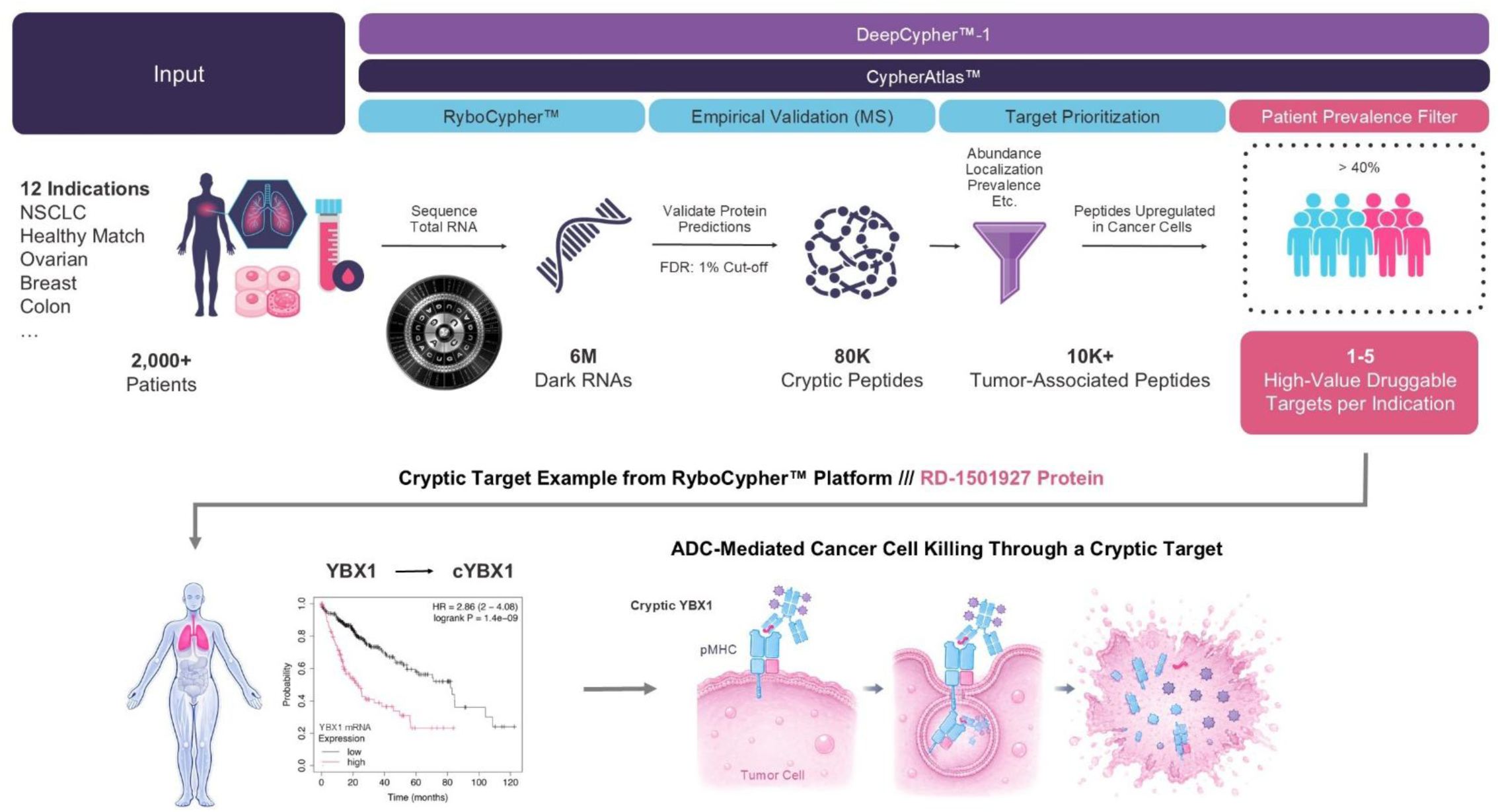

## Introduction

Few forces have reshaped modern medicine as decisively as the discovery of previously hidden biology. Releasing the immune system’s molecular “brakes” produced durable, and even curative responses in once-untreatable cancers. This work was recognized by the 2018 Nobel Prize^1,2^, while CAR T cells cured refractory leukemias and lymphomas^3^, and checkpoint blockade gave metastatic melanoma long-term survivors^4^. The lesson of the last two decades is unambiguous: each time the field uncovers a genuinely new layer of biology, a wave of transformative of first-in-class medicines follows.

Yet the catalog of actionable targets remains startlingly narrow. Fewer than 700 human proteins are engaged by any approved drug, and only a small minority of cancer driver genes are addressed by current therapies^5,6^, a limited repertoire that drives resistance and relapse, especially in solid tumors^7^. Even the best targeted immunotherapies reach only a fraction of patients: ∼15-20% of breast tumors are HER2-positive^8^, and PD-L1 is expressed at therapeutically relevant levels in only a subset of tumors^9^. The central constraint is specificity: because cancer arises from healthy cells, truly cancer-specific targets are exceedingly rare^10^. Next-generation sequencing revealed extensive molecular differences between tumor and normal tissue, yet, anchored to well-annotated RNA when inferring proteins, the field has systematically excluded much of the molecular output unique to malignant cells, for methodological rather than biological reasons.

Proteogenomic evidence indicates that tumors produce a far richer repertoire than reference annotations capture^11–14^ including non-canonical and highly structured RNAs that yield stable protein products detectable by mass spectrometry^14,15^. These poorly characterized “dark” RNAs can encode cryptic peptides, and tumors have been shown to generate protein products from transcriptional and translational events that are absent from standard annotations^12,16,17^. Recent reports of cryptic protein expression have provided evidence for thousands of novel peptides, but the majority of these are expressed from relatively short predicted open reading frames (ORF) or generate small peptides, which may limit the therapeutic utility of these cryptic protein products^18^. Although highly desired, novel resident membrane proteins bearing molecularly distinct extracellular domains have been elusive, as these represent an attractive class of therapeutic targets for conventional biologics. However, cryptic intracellular proteins that would otherwise remain inaccessible to immunotherapy become targetable through antigen processing, whereby proteins are processed into short peptides displayed on the cell surface in complex with human leukocyte antigen (peptide-MHC, or pMHC)^19^. Together, the discovery of cancer-associated cryptic proteins of these classes would substantially expand the repertoire of actionable tumor antigens, enabling both direct targeting of novel cell-surface proteins and pMHC-directed immunotherapies. This creates opportunities for a broad range of therapeutic modalities, including conventional antibodies, antibody-drug conjugates (ADCs), TCR-mimic antibodies (TCRm), and TCRm-based bispecific T-cell engagers (TCEs). Advances in antibody engineering now enable discrimination of single amino acid differences in displayed peptides, mitigating the specificity and off-tumor toxicity concerns that constrained earlier approaches^20,21^.

Here we introduce RyboCypher™, a novel integrated RNA-sequencing and AI-assisted proteogenomics platform that systematically maps unannotated peptides to the dark transcriptome generated by RyboCypher. RyboCypher identifies novel full-length RNAs at single-base resolution and integrates computational prediction with empirical proteomic validation to construct CypherAtlas™, a comprehensive proteogenomic atlas of the dark proteome. We provide empirical evidence for thousands of novel peptides and full-length proteins expressed from the RyboCypher-derived dark transcriptome; a cryptic proteome of intracellular proteins (transcription factors, E3 ligases) and membrane proteins (adhesion molecules, transporters, receptors), with many fragments detectable in the immunopeptidome. Applying a multimodal prioritization framework, we nominate highly conserved, cancer-associated pMHC candidates and full-length cryptic transmembrane targets for therapeutic development. As proof of concept, we advance one pMHC target candidate to TCRm antibodies that discriminate non-small cell lung adenocarcinoma (LUAD) cell lines with sub-nanomolar affinity, mediate potent ADC-induced tumor cell killing, and bind across three related HLA haplotypes, expanding the treatable patient population. Together, these results establish a new layer of biology for therapeutic targeting and a streamlined route to first-in-class biologics.

## Results

### Discovery of a Dark Transcriptome with RyboCypher™

RyboCypher resolves highly structured, full-length RNA species at single-base resolution (Graphical Abstract). A large fraction of the molecules it captures are invisible to conventional library preparation and sequencing methods and therefore, remain unannotated in literature. To systematically map the dark transcriptome, we applied RyboCypher to LUAD cell lines (A549, H1650), 8 LUAD tumors, and 3 colorectal cancer (CRC) tumor sections, with three matched healthy adjacent lung samples as reference. Across all samples, non-canonical transcripts were consolidated by locus-based aggregation, isoform collapsing, and reproducibility filtering. In total, we detected 6,605,037 unique “dark” RNA loci (Fig. 1a) and 8,308,621 unique isoforms across the cell lines and tissue samples analyzed. Our platform’s protein prediction module generated 16,103,687 cryptic protein predictions, distributed across dark RNA obtained from CRC tumors (1,024,598 loci; 15.5%), LUAD tumors (3,598,701; 54.5%), matched healthy LUAD (2,287,241; 34.6%), and LUAD cancer cell lines (105,891; 1.6%) (Supplementary Fig. 1a). We find that these transcriptional programs are broadly recurrent, rather than cancer-type-restricted (Supplementary Fig. 1b). The dark RNA transcripts sequenced by RyboCypher arise from strikingly consistent genomic regions: mostly from exon-intron junctions (40.2%), then intronic (37.2%), antisense (9.6%), exonic (9.2%), and intergenic (3.8%) loci (Supplementary Fig. 1c). This stability points to structured RNA processing rather than stochastic transcription. When mapped against annotated genes, the identified transcripts span 13,309 Ensembl gene IDs corresponding to 11,002 unique gene names (Supplementary Fig. 1d), representing a reproducible non-canonical layer, and providing a stable substrate for protein inference.

**Fig. 1.**
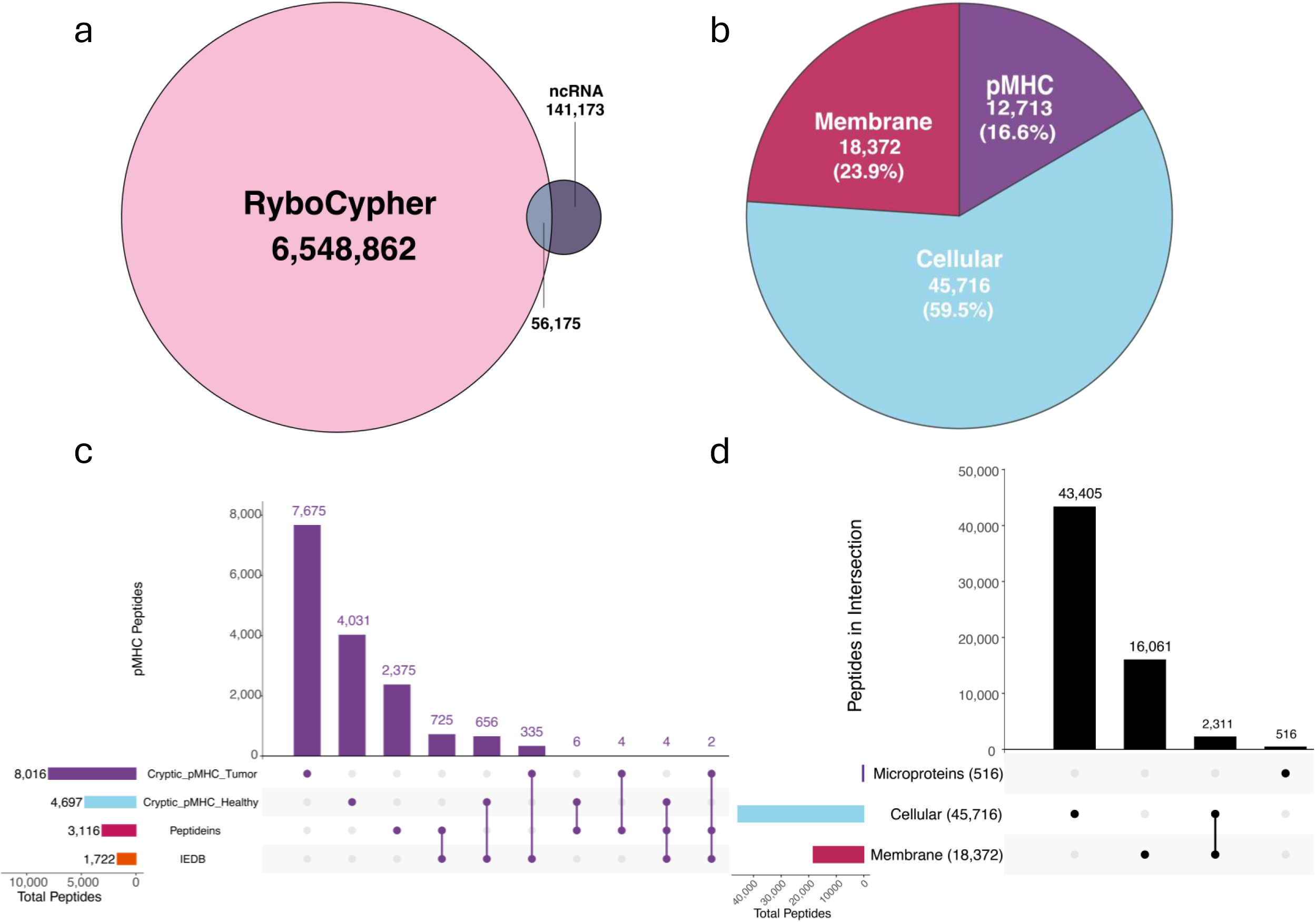
RyboCypher as a discovery engine for dark RNA and their encoded cryptic proteins. (Graphical Abstract) Schematic of the RyboCypher and downstream platform workflow. Two cancer cell lines along with patient tumor and healthy tissue samples (14 patients across NSCLC and CRC) are processed through RyboCypher to generate a dark transcriptome reference (∼6M dark RNAs). Proteogenomic searching within CypherAtlas identifies ∼80K cryptic peptides and application of an indication filter yields ∼10,000 cancer-associated peptides. 1-5 prioritized targets are selected for antibody development; an exemplary cryptic cancer-restricted peptide expressed from the YBX1 locus is targeted by a TCRm antibody formatted as ADCs. **(a)** Venn diagram comparing RyboCypher-derived non-canonical (ncRNA) loci (6,548,862) relative to known ncRNA databases; 56,175 loci are shared with ncRNA databases and 141,173 are present only in ncRNA databases, demonstrating that the vast majority of the RyboCypher-derived transcriptome is novel relative to existing ncRNA annotations. **(b)** Pie chart showing relative composition of RyboCypher-derived cryptic peptides identified by CypherAtlas: searching of raw MS data files obtained from samples in-house, and from publicly available proteomics datasets derived from over 2,000 patient samples revealed thousands of unannotated peptides in cellular proteomics (45,716; 59.5%), membrane proteomics preparations (18,372; 23.9%), and displayed peptides (pMHC) in the immunopeptidome (12,713; 16.5%). **(c)** Upset plot: CypherAtlas cryptic pMHC intersection with HLA-I Peptideins (Deutsch et al. 2026, Table S7) and the Immune Epitope Database and Analysis Resource (IEDB)^22^. CypherAtlas cryptic tumor-associated (pMHC_Tumor; 8,011), CypherAtlas cryptic healthy (pMHC_Healthy; 4,697), Peptideins (3,116), IEDB (1,722). The largest intersection bar (7,675) represents tumor-associated peptides identified by CypherAtlas and are unique to this dataset. **(d)** Upset plot: CypherAtlas cryptic cellular peptides (Cellular) and membrane peptides (Membrane) intersected with the Microproteins database (Deutsch et al. 2026, Table S2); Membrane (18,372), Cellular (45,716), Microproteins (516). Largest bar (43,405) represents Cellular peptides unique to RyboDyn. The vast majority of cryptic peptides have no overlap with any existing protein catalogs.

The samples sequenced by RyboCypher were supplemented with four matched healthy lung samples, 9 colon tumor samples, and 6 matched healthy colon samples and profiled for immunopeptidomics mass spectrometry. Mass spectra from each sample were searched against the protein predictions generated from RyboCypher dark RNA with subsequent targets prioritized using AI-assistance and empirical validation. CypherAtlas then applies disease-specific prioritization models to nominate cancer-associated therapeutic targets, including membrane proteins with targetable extracellular domains and pMHC-presented peptides conserved across tumors and common HLA haplotypes (e.g., HLA-A*02:01). Comparison against existing non-coding RNA databases revealed that the RyboCypher-derived dark transcriptome is overwhelmingly novel, with >97% of detected loci absent from current annotations. Initial proteogenomic interrogation within CypherAtlas provided empirical evidence that this dark transcriptome encodes a cryptic proteome spanning cellular, membrane, and immunopeptidome compartments (Fig. 1b), establishing a foundation for systematic therapeutic target discovery.

### Defining a Cryptic Human Proteome with CypherAtlas™

To determine whether the predicted cryptic proteins are translated, we constructed CypherAtlas, a proteogenomic search database integrating the Ensembl reference proteome, the nuORF database, and RyboCypher-derived protein predictions^14,17,23^. Peptide-spectrum matches were filtered at 1% FDR and required at least one amino acid mismatch relative to any reference or nuORF sequence, providing stringent evidence for bona fide cryptic translation products. We searched internal and public immunopeptidomics together with CPTAC cellular and membrane proteomics datasets, identifying 45,716 Cellular-MS, 18,372 Membrane-MS, and 12,713 pMHC-MS unique cryptic peptides across 12 different cancer types (Fig. 1b, 2a,c, Supplementary Tables 1-2)^14,23^. Comparison against catalogued microproteins and peptideins^18^ demonstrated that the overwhelming majority of these peptides have not been previously reported establishing CypherAtlas as a substantial expansion of the known human proteome (Fig. 1c, d). Specifically, direct intersection of RyboCypher cryptic pMHC against the HLA-I Peptideins catalog and of Cellular-MS and Membrane-MS peptides against the Microproteins database confirmed that the vast majority of RyboCypher peptides are unique to this dataset (Fig. 1c,d).

The cryptic proteome exhibited hallmarks consistent with biological and therapeutic relevance. Data derived from 10 different oncology indications from the CPTAC repository provided thousands of peptides matching CypherAtlas predicted proteins from both cellular and membrane protocols (Fig. 2a). The scale of peptides detected from each indication reflected cohort size and mass spectrometry analysis depth, with the largest peptide sets in Gastric (14,472) and LUAD (13,857) cellular proteomics datasets. To validate that our reanalysis is consistent with the published CPTAC results, we compared canonical protein fold-change values and observed strong concordance across all cohorts (Pearson r = 0.69-0.96; Supplementary Fig. 2b), indicating no systematic bias in our data processing pipeline.

**Fig. 2.**
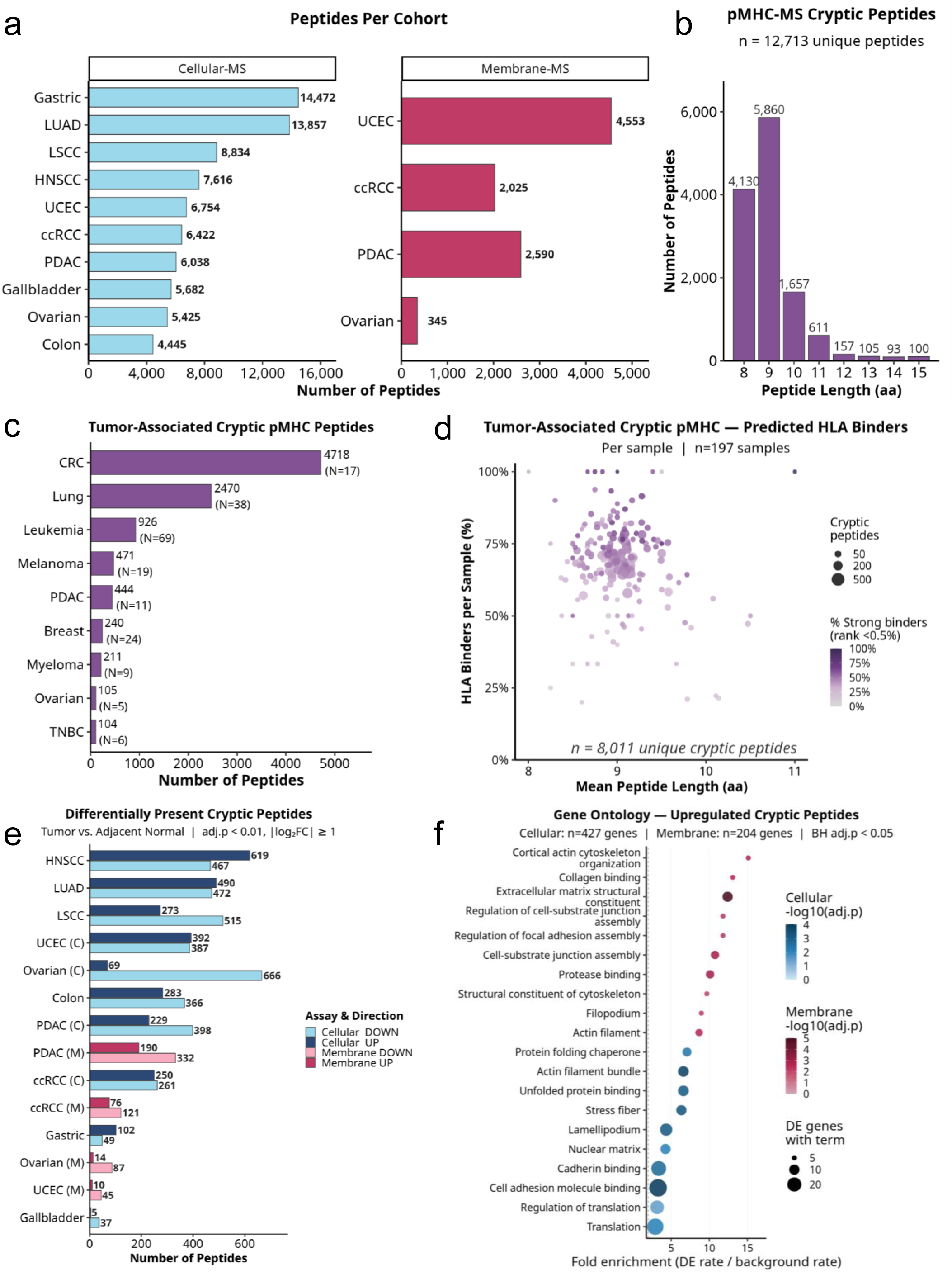
Discovery and initial characterization of a cryptic human proteome. **(a)** Unique cryptic peptides per cancer cohort by modality. Left: Cellular-MS cohorts (gastric [stomach] cancer (GS), lung squamous cell carcinoma (LSCC), lung adenocarcinoma (LUAD), head and neck squamous cell carcinoma (HNSCC), uterine corpus endometrial carcinoma (UCEC), clear cell renal cell carcinoma (ccRCC), pancreatic ductal adenocarcinoma (PDAC), gallbladder carcinoma (GB), Ovarian, Colon (CO)). Right: Membrane-MS cohorts (UCEC, ccRCC, PDAC, Ovarian). **(b)** pMHC-MS cryptic peptide length distribution (n = 12,713 unique peptides), showing the expected 8-11 aa HLA class I distribution with peak length at 9 aa (5,859 peptides). **(c)** Tumor-associated cryptic pMHC peptides per cancer type, stratified by tumor-specificity filter (All, Tumor-Associated, detected in >=2 tumors [TA]). Sample sizes (N) indicated per cohort. **(d)** Tumor-associated cryptic pMHC - predicted HLA binders. Per sample (n = 197 samples; n = 8,011 unique cryptic peptides). Dot size = peptide count; color = % strong binders (MHCflurry rank <0.5%); x-axis = mean peptide length. **(e)** Differentially present cryptic peptides per cohort (Tumor vs. Adjacent Normal; adj. p < 0.01, |log2FC| >= 1), Cellular-MS (blue) and Membrane-MS (pink). UP and DOWN counts shown per cancer type. **(f)** Gene Ontology - upregulated cryptic peptides. Cellular-MS (n = 427 genes) and Membrane-MS (n = 204 genes) cohorts (BH adj. p < 0.05). Dot size = number of DE genes with term; color encodes-log10(adj. p).

Evaluation of pMHC-derived cryptic peptides revealed the expected HLA class I length distribution, peaking at 9 amino acids (5,859 peptides; Fig. 2b), consistent with canonical antigen presentation. Tumor-associated cryptic pMHC peptides were identified across multiple cancer types, with the largest sets in CRC (4,718; N = 17) and Lung (2,470; N = 38) (Fig. 2c). To assess the immunogenic potential of these tumor-associated cryptic peptides, we predicted HLA class I binding affinity for each peptide against a panel of 19 common HLA-A, -B, and -C alleles using MHCflurry^58^. Across 197 patient samples comprising 8,011 unique cryptic peptides, we found that 55-80% of peptides detected per sample were predicted to bind at least one HLA class I allele with high affinity (percentile rank below 2%), with a substantial fraction qualifying as strong binders (percentile rank below 0.5%). These binding rates were consistent across samples regardless of the number of cryptic peptides detected or their mean peptide length, which ranged from 8 to 11 amino acids, consistent with canonical HLA class I ligand lengths (Fig. 2d). The high proportion of predicted binders across diverse HLA alleles suggests that tumor-associated cryptic peptides possess broad immunogenic potential suitable for therapeutic targeting across genetically diverse patient populations.

Tandem mass tags (TMT) enabled quantitative differential expression of cryptic proteins in parallel with canonical proteins from the same samples (Supplementary Fig. 2). Differential abundance analysis using MSstatsTMT^24^ identified numerous tumor-enriched cryptic peptides across both cellular and membrane proteomes (Fig. 2e, Supplementary Fig. 3). The loci encoding these cryptic peptides are enriched in protein classes the industry has long pursued for therapeutic intervention. Cellular proteomics revealed over-representation of cryptic deubiquitinases, E3 ligases, and transcription factors (Fig. 2f, Supplementary Fig. 4); whereas membrane proteomics extended this to cell adhesion molecules, transporters, and receptors, among the most coveted target classes in drug discovery, now surfacing in cryptic form.

### Prioritization of Cryptic Therapeutic Targets with AI models

Nominating therapeutic targets from the cryptic proteome requires integrating evidence across the transcript, protein, and patient dimensions at scale. To this end, we are developing DarkCypher-1, a multimodal deep-learning model that jointly encodes dark RNA sequences, predicted protein sequences, and structured biological metadata through modality-specific encoders, links transcript and protein representations using cross-attention, and conditions predictions on sample context to generate embeddings for downstream therapeutic prioritization (Fig. 3a). CypherAtlas provides the proprietary experimental data and metadata that serve as its training substrate; within this architecture, a protein language model such as ESM-2 can serve as the protein encoder.

**Fig. 3.**
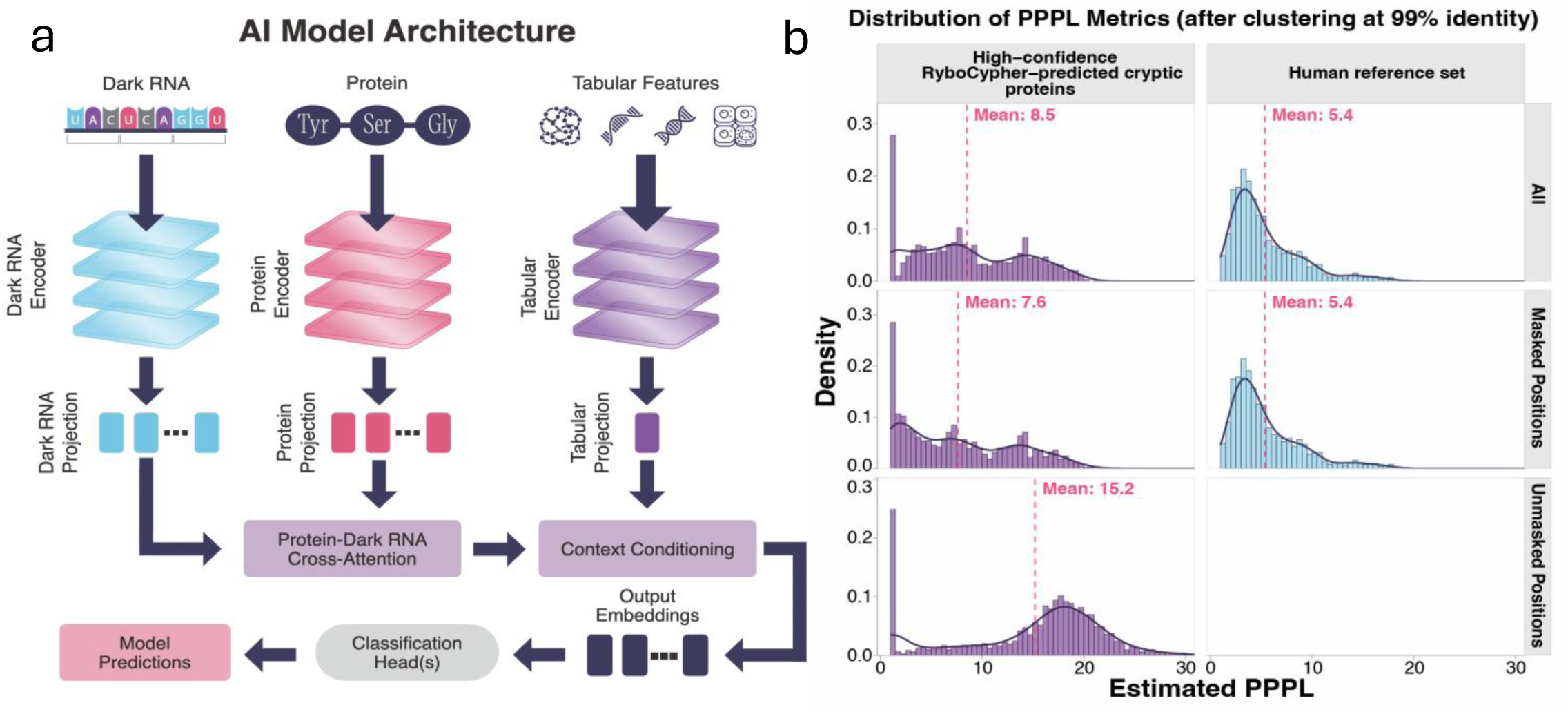
CypherAtlas integrates omics datasets with metadata and enables AI-guided cryptic target prioritization. **(a)** Schematic of DarkCypher-1, a multi-modal deep-learning architecture in development that predicts target expression in patient samples. Modality-specific encoders embed the candidate dark-RNA transcript, its predicted protein, and structured sample-level metadata (protein identity, expression, patient haplotype, and cancer type); cross-attention links the transcript and protein representations, and a context-conditioning step incorporates the sample metadata before a classification head outputs the prediction. Because no existing model addresses RyboCypher’s drug-target modality, the architecture is designed to be trained directly on RyboDyn’s proprietary data to improve sensitivity, reduce false discovery, and accelerate target selection, with additional data expected to enable population-and cancer-specific target discovery and in-silico screening. **(b)** Distributions of direct PPPL for high-confidence RyboCypher-predicted cryptic proteins (n = 3,192) and the human reference set (n = 1,017), after MMseqs2 clustering at 99% identity. Top row: whole-sequence PPPL; middle and bottom rows decompose PPPL into masked-position (reference-covered) and unmasked-position (novel) geometric means. Dashed lines mark per-facet means.

The ∼8.3 million dark RNA isoforms identified by RyboCypher™ give rise to approximately 16 million candidate open reading frames (ORFs), creating a vast search space in which only a subset are expected to encode bona fide proteins. Because cryptic proteins lack experimental annotation, we asked whether ESM recognized them as plausible biological proteins rather than statistical artifacts. We first verified that ESM-2 pseudo-perplexity (PPPL) is calibrated to protein plausibility across a panel of sequence perturbations, and that a fast single-pass approximation (direct PPPL) closely reproduces the exact metric for cohort-scale scoring (Methods; Supplementary Fig. S5-S8). Using this PPPL metric, we quantified how closely each predicted cryptic protein conformed to the statistical properties of naturally occurring proteins. High-confidence RyboCypher predictions (n = 3,192; MMseqs2 clustered at 99% identity)^25^ produced whole-sequence PPPL distributions that overlapped substantially with those of curated human reference proteins at the low-PPPL end but were systematically shifted toward higher PPPL, with a heavier right tail (mean 8.5 versus 5.4; Fig. 3b). Importantly, reference-covered (masked) regions were nearly indistinguishable from canonical proteins (mean 7.6 versus 5.4), whereas elevated PPPL was confined almost exclusively to the novel out-of-reference (unmasked) regions (mean 13.2). These findings indicate that cryptic proteins are systematically less plausible than canonical proteins, occupying an intermediate plausibility band; clearly separated from random or shuffled sequences yet distinguishable from the canonical proteome, with this elevation driven almost entirely by their genuinely novel, out-of-reference regions. Because the majority of cryptic proteins consist of masked and unmasked regions, we next resolved PPPL to individual residues, labeling each position as reference-supported or novel by reference k-mer encoding. Nearly all high-confidence candidates (3,129 of 3,192; 98.0%) were mosaics of both, and in 88.7% the novel sub-regions were the least plausible part of the protein, elevated well above their reference-supported regions. This positional structure is directly actionable: residues that are simultaneously absent from existing reference annotations while remaining strongly supported as cryptic protein sequence, making them attractive candidates for selective therapeutic targeting.

### Prioritization of Actionable Cryptic Cell-Surface Targets

To prioritize cryptic ORF-derived cell-surface targets from multi-cancer mass spectrometry data, we implemented a composite scoring framework within CypherAtlas that integrates differential expression evidence across cohorts with orthogonal membrane topology predictions. Candidate cryptic peptides upregulated in tumor versus healthy tissue proteomes were first filtered to those arising from proteins with predicted transmembrane architecture, as determined by TMbed^26^ run locally. Select top candidate proteins were further evaluated using DeepTMHMM^27^ to confirm topology, resolve domain orientation, and visualize per-residue architecture^26–28^. A pan-cancer membrane score combined the maximum observed log2 fold change across cohorts, the number of cancer types with significant upregulation, the DeepLoc 2.0 cell membrane probability, and a concordance bonus when both TMbed and DeepTMHMM independently confirmed transmembrane topology (Fig. 4a). Together, these orthogonal features prioritize cryptic proteins that are reproducibly enriched in tumors while exhibiting high-confidence membrane localization and topology consistent with cell-surface accessibility. The top-ranked candidates (peptide differential abundance shown in Supplementary Fig. 9) span diverse membrane topologies, including type-I single-pass proteins with signal peptides, multi-pass polytopic membrane proteins, and proteins with extracellular domains accessible to antibody engagement. The four highest-scoring proteins were selected for detailed topology visualization (Fig. 4b), illustrating that membrane localization alone is insufficient for therapeutic prioritization. Rather, the detected cryptic peptide must be physically accessible from outside the cell while residing within a tumor-associated cryptic sequence.

**Fig. 4.**
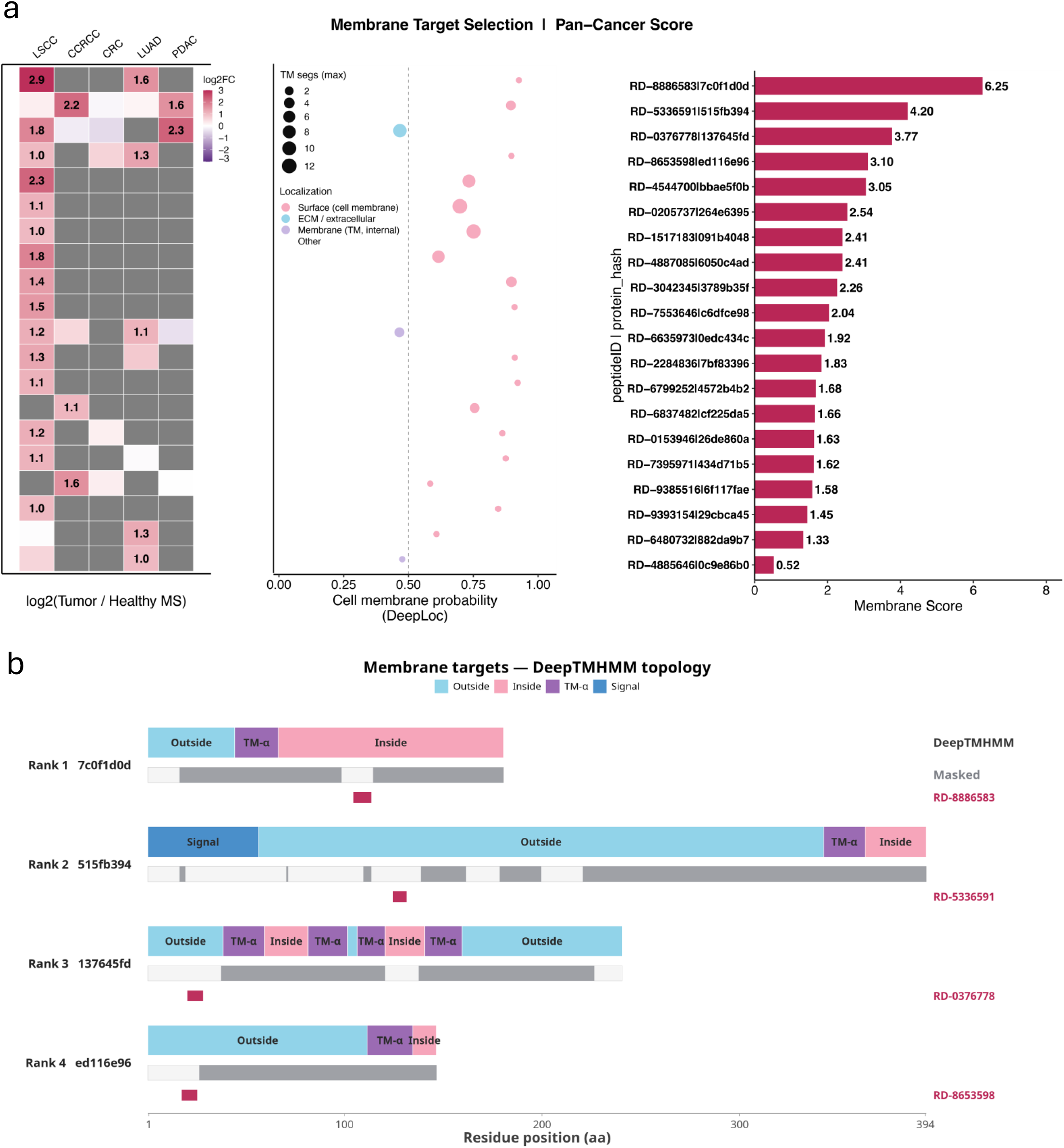
Selection of actionable membrane targets from the cryptic cancer proteome. **(a)** Membrane Target Selection: Pan-Cancer Score. The heatmap (left panel) shows log2(Tumor/Healthy MS intensity) per cancer cohort (ccRCC, CRC, LSCC, LUAD, PDAC); grey cells indicate the peptide was not detected or did not meet the fold-change threshold in that cohort. The dot plot (middle panel) shows DeepLoc 2.0 cell membrane probability for each candidate; dot size encodes the maximum predicted TM segment count across both tools; dot color indicates DeepLoc subcellular localization category (pink = surface/cell membrane; blue = ECM/extracellular; lavender = membrane, TM/internal; grey = other). Candidates are ordered by descending membrane score. Cryptic ORF-derived membrane candidates ranked by composite membrane score (horizontal bars, right panel). Each candidate is identified by its internal cryptic peptide identifier. Score components include maximum observed tumor/healthy, log2 fold change across all cohorts, number of cancer types with upregulation (log2FC ≥ 1.0), DeepLoc 2.0 cell membrane probability, transmembrane segment count (TMbed and DeepTMHMM consensus), and a concordance bonus for dual-tool TM agreement. **(b)** DeepTMHMM Topology: Top 4 Membrane Candidates. DeepTMHMM topology for the four highest-scoring cryptic ORF candidates. Bars are scaled to protein length; colors indicate per-residue domain assignment: Outside (light blue), Inside (pink), TM-α helix (purple), Signal peptide (dark blue). Grey track shows residues masked by reference k-mer coverage (see Methods). Red band marks the position of the cryptic MS-detected peptide within the ORF. The four candidates exemplify distinct membrane topologies supporting surface accessibility: Rank 1 encodes a single-pass protein with the detected peptide located in the intracellular domain, flanked by a single TM helix; Rank 2 carries an N-terminal signal peptide followed by a large extracellular domain with the detected peptide positioned in the outside region, terminated by a single C-terminal TM helix, a classical type-I surface topology; Rank 3 is a multi-pass polytopic membrane protein with the detected peptide in an extracellular loop between TM helices; and Rank 4 is a single-pass protein with the detected peptide in the extracellular outside region proximal to the TM helix.

The masked track in Fig. 4b is central to cell-surface target selection. Grey residues denote regions covered by canonical reference k-mers (Methods), representing sequence shared with the known human proteome and therefore carrying potential cross-reactivity risk against normal tissues. In contrast, unmasked residues correspond to genuinely cryptic, non-canonical sequence that confers tumor specificity which is the same masked/unmasked distinction that underlies the PPPL analysis in Fig. 3b. An actionable membrane target must place its MS-detected cryptic peptide (red band) within both an unmasked (non-canonical) region and an extracellular or membrane-exposed domain, thereby combining tumor-associated sequence novelty with physical accessibility for antibody engagement. The top-ranked candidates satisfy both requirements jointly: RD-5336591 adopts a classical type-I topology, with an N-terminal signal peptide, a large extracellular domain carrying the detected cryptic peptide, and a single C-terminal transmembrane helix; RD-0376778 is a polytopic protein displaying its cryptic peptide in an extracellular loop; and RD-8653598 exposes its cryptic peptide in the outside region proximal to the transmembrane helix (Fig. 4b). Together, these results demonstrate that CypherAtlas systematically prioritizes cryptic cell-surface proteins possessing both extracellular accessibility and tumor-associated sequence novelty, representing attractive candidates for antibody-based therapeutics.

### Discovery of Recurrent Cryptic pMHC Targets Across Human Cancers

To evaluate whether cryptic pMHC targets recur across independent cancer types, we re-searched published immunopeptidomics datasets using RyboCypher-derived protein predictions together with our in-house patient datasets. Across all indications, CypherAtlas recovered thousands of cryptic pMHCs, a substantial fraction of which were absent from healthy tissues and were recurrent across independent tumors (Fig. 5a). Colorectal cancer yielded the largest cryptic immunopeptidome (7,361 total; 4,718 tumor-associated; 3,501 recurrent), followed by lung cancer (5,511; 2,470; 1,918), while leukemia, melanoma, PDAC, myeloma, breast, TNBC, and ovarian cancers each contributed hundreds of recurrent tumor-associated cryptic pMHCs. The higher discovery rate in lung and CRC was expected, as the underlying dark RNA reference was generated almost exclusively from RyboCypher sequencing of colorectal and lung tumors, providing substantially greater discovery depth for these indications.

**Fig. 5.**
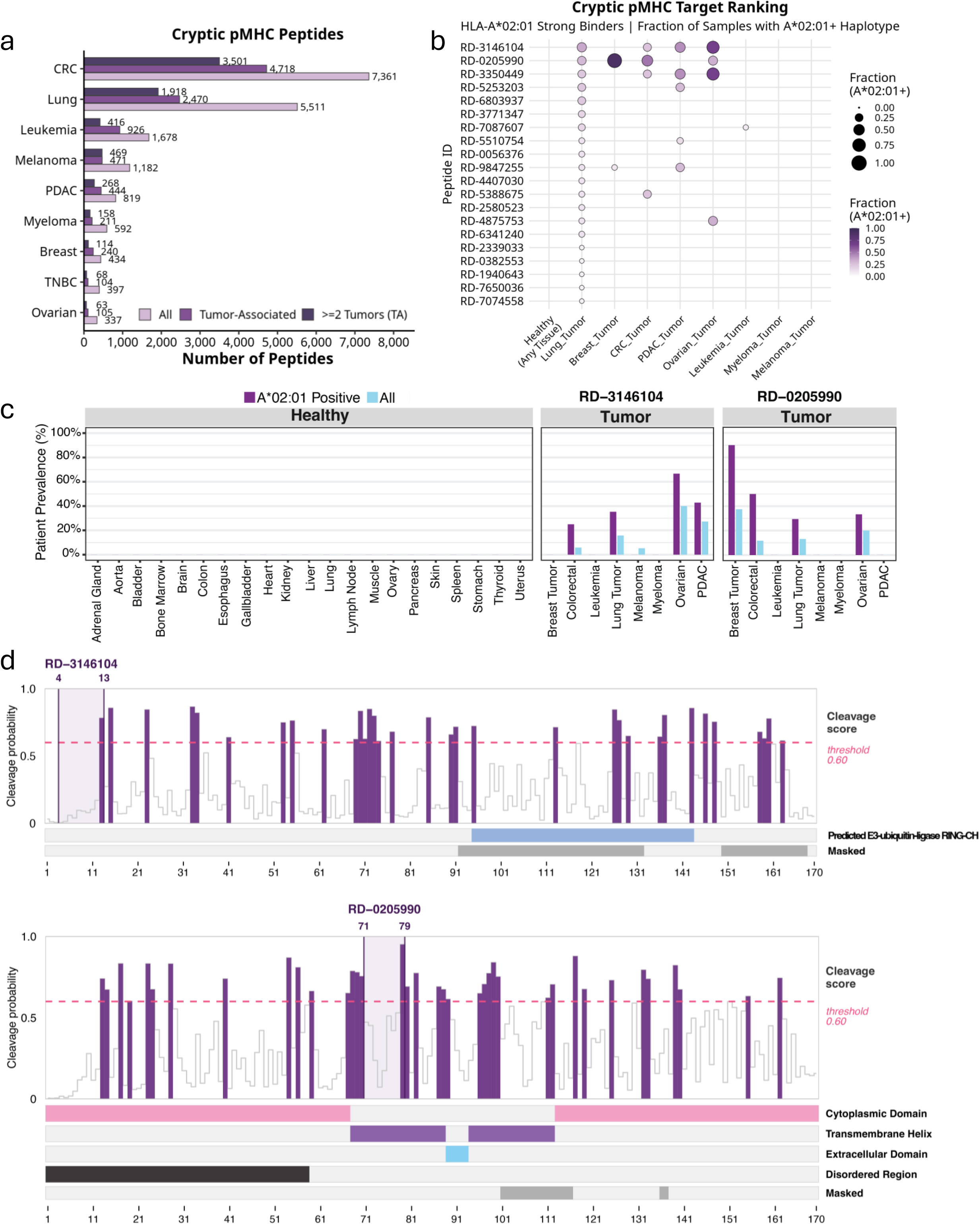
Selection of actionable targets from the cryptic cancer immunopeptidome. **(a)** Number of DarkRNA-encoded cryptic peptides detected by pMHC-MS across cancer types, stratified by tumor-association filter. Light purple: all cryptic peptides containing at least one mismatch from the reference proteome. Medium purple: tumor-associated subset not detected in any healthy tissue (HLA Healthy Ligand Atlas^30^ or donor-matched healthy pMHC-MS). Dark purple: tumor-associated peptides detected in two or more independent tumor samples. CRC: 7,361 / 4,718 / 3,501; Lung: 5,511 / 2,470 / 1,918; Leukemia: 1,678 / 926 / 416; Melanoma: 1,182 / 471 / 469; PDAC: 819 / 444 / 268; Myeloma: 592 / 211 / 158; Breast: 434 / 240 / 114; TNBC: 397 / 104 / 68; Ovarian: 337 / 105 / 63. **(b)** Cryptic pMHC target ranking. Top 20 tumor-associated DarkRNA cryptic peptides ranked by HLA-A02:01 strong binding (MHCflurry rank <0.5%) and fraction of HLA-A02:01-positive samples with detected presentation by pMHC-MS across cancer types. Dot size encodes fraction of HLA-A*02:01-positive patients in each indication; dot color intensity encodes the same metric. RD-3146104 and RD-0205990 rank first and second, respectively, with presentation confirmed across multiple independent cancer types. **(c)** Patient prevalence of RD-3146104 and RD-0205990 across healthy tissues and tumor types. Purple: HLA-A02:01-positive patients; blue: all patients. Left panels show absence of detection across 19 healthy tissue types from the HLA Healthy Ligand Atlas; right panels show tumor-restricted detection in colorectal, lung, leukemia, myeloma, ovarian, and PDAC cohorts. Tumor exclusivity confirmed against the HLA Healthy Ligand Atlas and donor-matched healthy pMHC-MS datasets. **(d) (top)** Proteasomal cleavage prediction for RD-3146104 (193 aa ORF; masked: 35.8% canonical sequence similarity); epitope spans residues 4-13 near the ORF N-terminus, upstream of the predicted E3 ubiquitin-ligase RING-CH domain. Single high-confidence cleavage predicted at the C-terminal residue with no internal cleavage above threshold. **(bottom)** Proteasomal cleavage prediction for RD-0205990 (191 aa ORF; masked: 9.4% canonical sequence similarity); epitope spans residues 71-79 at the cytoplasmic-transmembrane boundary. Strong C-terminal cleavage predicted at position 79 with no internal cleavage.

Ranking tumor-exclusive strong binders by cross-cancer recurrence revealed a compact set of shared therapeutic antigens (Fig. 5b). RD-3146104 and RD-0205990 emerged early as two high-priority candidates, with presentation independently confirmed across multiple cancer types in HLA-A*02:01-positive patients. RD-3146104 originates from a cryptic ORF encoding an E3 ubiquitin ligase, whereas RD-0205990 derives from a focal adhesion protein; both peptides were detected exclusively in tumor immunopeptidomes despite repeated observation across independent patient cohorts.

The leading candidates also demonstrated exceptional patient coverage while remaining undetectable in healthy tissues (Fig. 5c). RD-3146104 was identified in colorectal, lung, ovarian, and pancreatic cancers, reaching nearly 65% prevalence among HLA-A02:01-positive ovarian tumors. RD-0205990 was detected across breast, colorectal, lung, ovarian, and pancreatic cancers and reached approximately 90% prevalence among HLA-A02:01-positive breast tumors. Importantly, neither peptide was detected across 19 surveyed healthy tissue types nor in donor-matched healthy immunopeptidomes. This degree of prevalence is remarkable; for comparison, only ∼15–20% of breast cancers overexpress HER2, whereas RD-0205990 was identified in approximately 40% of patients prior to stratifying by HLA haplotype, and in 90% of HLA-A*02:01-positive HER2-negative breast tumors. Together, these findings demonstrate that CypherAtlas identifies shared, tumor-restricted cryptic antigens with the prevalence and specificity required for broad therapeutic development.

### Prioritization of Cryptic pMHC Targets for Immunotherapy

While cryptic cell-surface proteins are directly accessible to antibody-based therapeutics, the majority of cryptic proteins reside intracellularly and therefore require an alternative route to therapeutic engagement. Antigen processing provides precisely one such mechanism, converting intracellular proteins into peptide-MHC (pMHC) complexes displayed on the cell surface. We therefore developed a complementary prioritization framework within CypherAtlas to identify cryptic pMHC targets with the greatest potential for immunotherapy.

To convert tens of thousands of translated cryptic peptides into a tractable therapeutic shortlist, CypherAtlas applies a sequential oncology filter across 154 patient samples spanning 10 different publicly available datasets (Supplementary Table 2). Candidate peptides must derive from biologically plausible cryptic proteins, be novel relative to the canonical proteome, absent from healthy tissues, recurrent across independent tumors, and presented by clinically prevalent HLA alleles (Supplementary Fig. 10). Candidates are further prioritized by integrating evidence for clinical actionability, including parent-locus association with patient outcome and predicted proteasomal processing (Fig. 5d). This framework reduces 12,713 unique candidate peptides to 8,011 tumor-associated peptides (not observed in any healthy sample or reference database), of which 5,301 were observed in at least 2 tumor samples, and 226 of these were highly conserved (>20% of samples). Following filtering for predicted binding to HLA-A*02:01, the list was further narrowed to 60 potential therapeutic target candidates. Synthetic versions of these top 60, cancer-restricted peptides were used for spectral validation and for determination of physiochemical properties, confirming 57 of 60 (95%).

Clinical prioritization relies on two complementary criteria. First, expression of the parent locus expressing the dark-RNA is evaluated for association with patient outcome; for example, high expression of the locus encoding RD-1501927 is associated with significantly reduced overall survival (TCGA, n = 105; Fig. 6a), indicating that its activity is linked to tumor biology rather than incidental transcription. Second, candidate epitopes must be predicted to undergo productive proteasomal processing. RD-1501927, RD-0205990, and RD-3146104 each exhibit high-confidence C-terminal cleavage immediately adjacent to the epitope, with no predicted internal cleavage across the peptide window (Fig. 5d), supporting efficient generation and presentation by HLA class I molecules.

**Fig. 6.**
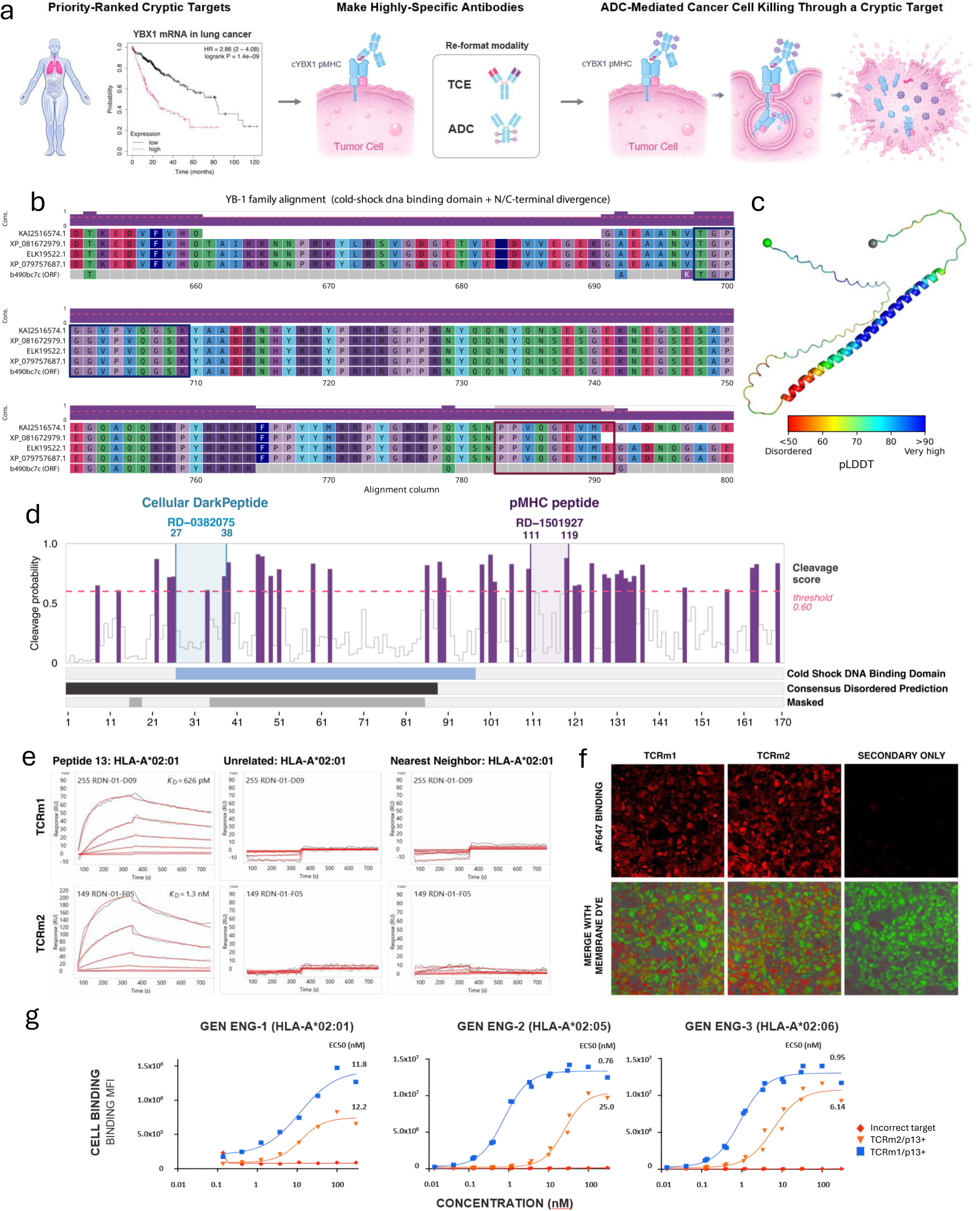
Targeting the dark proteome. **(a)** A cancer-restricted peptide-MHC encoded by the ybx1 locus is prioritized for targeting by TCRm antibodies. The locus is known to be deleterious to overall survival in lung and liver cancers when highly expressed at the RNA level (Kaplan-Meier plot for lung cancer shown; red curve = high expression, black curve = low expression). Cryptic proteins expressed in cancer cells are processed by the immunoproteasome, and displayed on the cell surface as pMHC targets. TCRm antibodies formatted as ADCs can kill cancer cells by internalization, causing for release of the toxic payload. **(b)** YB-1 family protein sequence alignment (cold-shock core + C-terminal divergence region) across human reference sequences and the RyboCypher-derived cryptic ORF (b490bc7c ORF). Alignment columns 660-800 shown; divergent C-terminal residues in the cryptic ORF confirm sequence novelty relative to canonical YBX1. **(c)** ESMFold2 structural model of the cYBX1 cryptic protein, colored by pLDDT confidence: blue (>90, very high), cyan (70-90, confident), yellow (50-70, low), orange (<50, very low). N-and C-termini labeled. **(d)** Proteasomal cleavage prediction for RD-1501927 (170 aa ORF; masked: 31.8% canonical sequence similarity); the epitope spans residues 111–119, outside the conserved DNA binding region referred to as the Cold Shock Domain (aa 27–87) in which the reference cellular peptide also falls. A high-confidence C-terminal cleavage site is predicted at position 119 with no internal cleavage within the epitope window. **(e)** SPR binding kinetics of TCRm1 and TCRm2 against Peptide 13 (RD-1501927) in HLA-A*02:01, an unrelated peptide, and the nearest-neighbor peptide. TCRm1 KD = 626 pM; TCRm2 KD = 1.3 nM. Neither clone shows measurable binding to unrelated or nearest-neighbor controls. **(f)** Fluorescence cell binding of TCRm1 and TCRm2 (AF647-labeled) on HLA-A*02:01-positive NSCLC cell lines. Top: AF647 signal (red). Bottom: merge with membrane dye (green). Secondary-only control shown. **(g)** Cell binding dose-response curves for TCRm1 (blue) and TCRm2 (orange) across three HLA haplotypes: HLA-A*02:01 (GEN ENG-1; EC50 11.8 nM and 12.2 nM), HLA-A*02:05 (GEN ENG-2; EC50 0.76 nM and 25.0 nM), and HLA-A*02:06 (GEN ENG-3; EC50 0.95 nM and 6.14 nM). Unrelated peptide control shows no antibody binding across all three haplotypes.

### Therapeutic Targeting of a Cryptic Protein Encoded by the YBX1 Locus

To demonstrate that cryptic pMHC targets identified by CypherAtlas can be therapeutically exploited, we selected RD-1501927 as an initial target for TCRm antibody generation (Fig. 6a,e-g). RD-1501927 is predicted to bind HLA-A*02:01 with moderate strength and HLA-A*02:05 and HLA-A*02:06 strongly. We identified dark transcripts in H1650 cells and LUAD tumor samples expressed from the ybx1 locus containing ORFs predicted to encode a cryptic protein containing the RD-1501927 peptide. Although the cryptic YBX1 protein (cYBX1) bears no significant homology to the human proteome, it aligns closely with predicted orthologs from the same locus in other species (Fig. 6b), it is predicted to fold into a full-length functional protein (Fig. 6c) containing disordered regions, and a highly conserved DNA binding region (Cold Shock Domain), suggesting evolutionary conservation across species, despite complete divergence from the canonical YBX1 protein.

To raise highly specific binders against RD-1501927, TCRm antibodies were selected by phage display from large human single-chain variable fragment (scFv) libraries. Following positive selection (bait: RD-1501927 in HLA-A*02:01) and negative selection (bait: unrelated peptide in HLA-A*02:01), HCDR3 regions from six scFv clones were advanced to affinity maturation, with more stringent negative selection using the nearest-neighbor peptide yielding clones of markedly improved specificity and affinity. Top clones were characterized by SPR against RD-1501927, an unrelated peptide, and the nearest-neighbor peptide, each displayed in HLA-A*02:01 (Fig. 6e). Thirty-one clones demonstrated high affinity and specificity for the target, with lead clones TCRm1 and TCRm2 binding at KD = 626 pM and 1.3 nM respectively. These antibodies exhibited no measurable binding to either an unrelated peptide or the nearest-neighbor peptide, despite the latter differing from RD-1501927 by only three amino acids. Four clones were converted to IgG for cell-binding studies on HLA-A*02:01-positive LUAD lines H2228 and H1650; both showed fluorescence above background, confirming presentation of RD-1501927 and specific TCRm binding on primary cell lines (Fig. 6f).

Anti-RD-1501927 TCRm antibodies were then profiled against engineered cells expressing pMHC polycistronic cassettes across three different HLA isoforms. Two clones productively bound RD-1501927, but not unrelated or nearest-neighbor peptides, when displayed in HLA-A*02:01, HLA-A*02:05, and HLA-A*02:06, with sub-nanomolar EC50 values in the more stable haplotypes (HLA-A*02:01: 11.8 and 12.2 nM; HLA-A*02:05: 0.76 and 25.0 nM; HLA-A*02:06: 0.95 and 6.14 nM for TCRm1 and TCRm2 respectively; Fig. 6g). Importantly, these results demonstrate that a single TCRm antibody can recognize the same cryptic pMHC across multiple closely related HLA-A*02 isoforms, effectively breaking haplotype restriction while still maintaining specificity to the displayed peptide. HLA-A*02:01 is present in ∼35-40% of Western European and North American populations, HLA-A*02:05 in 5-8%, and HLA-A*02:06 in 3-4%; a single antibody recognizing the target across all three isoforms raises the addressable population to over 45%. Initial testing of TCRm1 as an ADC (linker/payloads: lysosomal-cleavable linker, exatecan or eribulin) against HCC78 cells (HLA-A*02:05-positive) demonstrated cryptic pMHC target-specific cell killing *in vitro*, well above isotype controls (Supplementary Fig. 11). Together, these findings demonstrate that cryptic pMHCs discovered by RyboCypher and prioritized by CypherAtlas can be translated into highly specific therapeutic antibodies capable of recognizing endogenous tumor antigens across multiple clinically prevalent HLA haplotypes, providing a direct path from dark proteome discovery to first-in-class immunotherapies.

## Discussion

Here we present evidence indicating that the human proteome is substantially larger than currently appreciated. By integrating RyboCypher, CypherAtlas and DarkCypher-1, we map the RyboCypher-derived dark transcriptome onto a previously inaccessible dark proteome comprising tens of thousands of empirically identified novel peptides spanning intracellular, membrane, secreted and antigen-presented protein classes. These are not isolated artifacts but a reproducible and therapeutically accessible layer of biology that has been invisible to conventional genomics and proteomics. This proteome is distinct from recently described peptideins and microproteins, and absent from current reference annotations. As we extend this reference across additional healthy tissues and disease indications, and layer secretome, interactome and chemoproteome datasets onto continued cellular proteomics, membrane proteomics and immunopeptidomics, we anticipate that the catalog of therapeutically actionable cryptic proteins will grow in step, progressively widening the therapeutic modality and targetable landscape of the dark proteome.

Unlike previous proteogenomic studies that largely terminate at peptide discovery, our framework progresses through biological plausibility assessment, empirical proteomic validation, and modality-specific therapeutic prioritization. CypherAtlas integrates multimodal omics, patient metadata, and AI-guided prioritization to systematically nominate therapeutic targets across multiple drug modalities. By linking each cryptic target to cancer indication, patient prevalence, HLA genotype, expression, and clinical outcome at the earliest stages of discovery, CypherAtlas enables patient stratification before medicinal chemistry or antibody engineering has even begun. Because these richly annotated data are proprietary to RyboDyn and difficult to assemble elsewhere, CypherAtlas also constitutes a unique training substrate for AI models such as DarkCypher-1, which we are developing to prioritize cryptic targets and forecast their expression in new patient samples. Given that nearly 90% of oncology drugs entering clinical trials ultimately fail - often because the underlying biology or patient population is insufficiently understood - this target-first, patient-informed framework has the potential to fundamentally reduce the growing concentration of drug discovery efforts around a relatively small number of well-studied proteins.

Critically, the cryptic proteome emerging from this work is therapeutically addressable through multiple complementary modalities. Cryptic membrane proteins exposing extracellular domains provide opportunities for conventional antibodies, ADCs, radioligands, and other cell-surface targeting strategies, whereas intracellular cryptic proteins become accessible to small molecule intervention and through antigen processing and presentation, enabling TCR-mimic antibodies, bispecific T-cell engagers, and related immunotherapies. Rather than revealing a single class of therapeutic targets, the dark proteome substantially expands the spectrum of druggable biology across virtually every major therapeutic modality.

Among the therapeutic opportunities revealed here, we disclose cYBX1, a cryptic protein encoded from the YBX1 locus. Although the canonical YBX1 protein has long been recognized as a major oncogenic regulator, it has historically proven challenging to target directly. Remarkably, the cancer-associated dark RNA identified by RyboCypher gives rise to a previously unrecognized cryptic protein that, to our knowledge, generates a tumor-restricted pMHC displayed on the cell surface. This creates a new therapeutic handle on the YBX1 locus, enabling selective targeting where it matters while sparing healthy tissues. We demonstrate the generation of highly specific TCR-mimic antibodies against this cryptic pMHC target displayed on LUAD cell lines and engineered cells, with antibodies that recognize the correct peptide across multiple HLA haplotypes and thereby expand the treatable patient population. These antibodies now provide a foundation for *in vivo* evaluation of cryptic pMHC-directed therapeutics and illustrate the translational potential of targets emerging from the dark proteome. More broadly, these findings illustrate how the dark proteome can reveal tractable therapeutic opportunities within established cancer-associated loci that have historically remained difficult to drug.

Future studies will determine the biological function of cYBX1 and other cryptic proteins through genetic perturbation, pooled overexpression, and *in vivo* therapeutic evaluation. More broadly, systematic functional interrogation of cryptic ORFs should establish whether these proteins contribute directly to tumor fitness or represent lineage-restricted biomarkers that can be therapeutically exploited. Beyond oncology, the same framework should enable systematic discovery of cryptic cytokines, secreted proteins, enzyme families, and signaling pathways that remain inaccessible to current annotation, opening additional opportunities for conventional biologics and small-molecule discovery. Although this work focuses on cancer targeting, the healthy cryptic proteome discovered here represents a valuable resource. Tissue-restricted cryptic proteins may ultimately provide highly selective molecular addresses for targeted drug delivery, regenerative medicine, or tissue-specific imaging applications.

For decades, transformative advances in medicine have followed the discovery of previously unseen biology. If the annotated proteome defined the first era of targeted therapeutics, the dark proteome may define the next. By combining AI-guided discovery, proteogenomic validation, and therapeutic prioritization into a unified platform, RyboCypher, CypherAtlas and DarkCypher-1 provide not simply a catalog of previously unseen biology, but a new substrate for therapeutic discovery itself: one capable of systematically expanding the universe of first-in-class therapeutic targets for the patients they are designed to serve.

## Acknowledgements

The authors thank Dr. Steven P. Gygi, Professor of Cell Biology at Harvard Medical School, and the Gygi Laboratory for their partnership and for performing state-of-the-art proteomics analyses.

---

**Supplementary Fig. 1.**
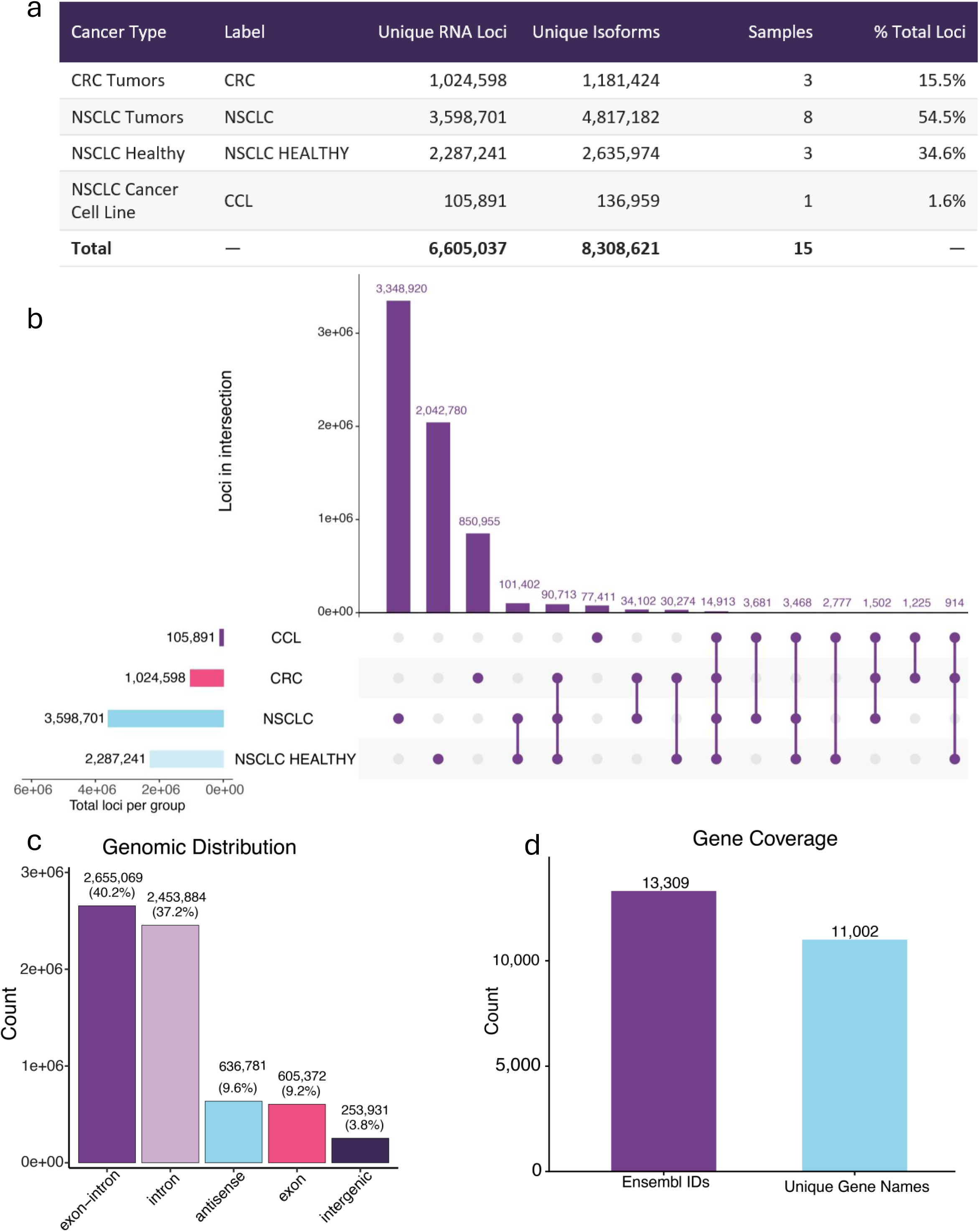
RyboCypher dark RNA characterization. **(a)** Summary table of unique RNA loci and isoforms per sample group: CRC Tumors (1,024,598 loci; 1,181,424 isoforms; n = 3; 15.5%), NSCLC Tumors (3,598,701; 4,817,182; n = 8; 54.5%), NSCLC Healthy (2,287,241; 2,635,974; n = 3; 34.6%), NSCLC Cancer Cell Line (105,891; 136,959; n = 1; 1.6%); total 6,605,037 loci / 8,308,621 isoforms across 15 samples. **(b)** Upset plot of loci intersections across sample groups. Largest single-group intersection: NSCLC tumors (3,348,920), followed by shared NSCLC/NSCLC-Healthy loci (2,042,780) and CRC-exclusive loci (850,955). **(c)** Genomic distribution of retained loci: exon-intron junctions (2,655,069; 40.2%), intronic (2,453,884; 37.2%), antisense (636,781; 9.6%), exonic (605,372; 9.2%), intergenic (253,931; 3.8%). **(d)** Gene coverage: 13,309 Ensembl IDs corresponding to 11,002 unique gene names.

**Supplementary Table 1.**
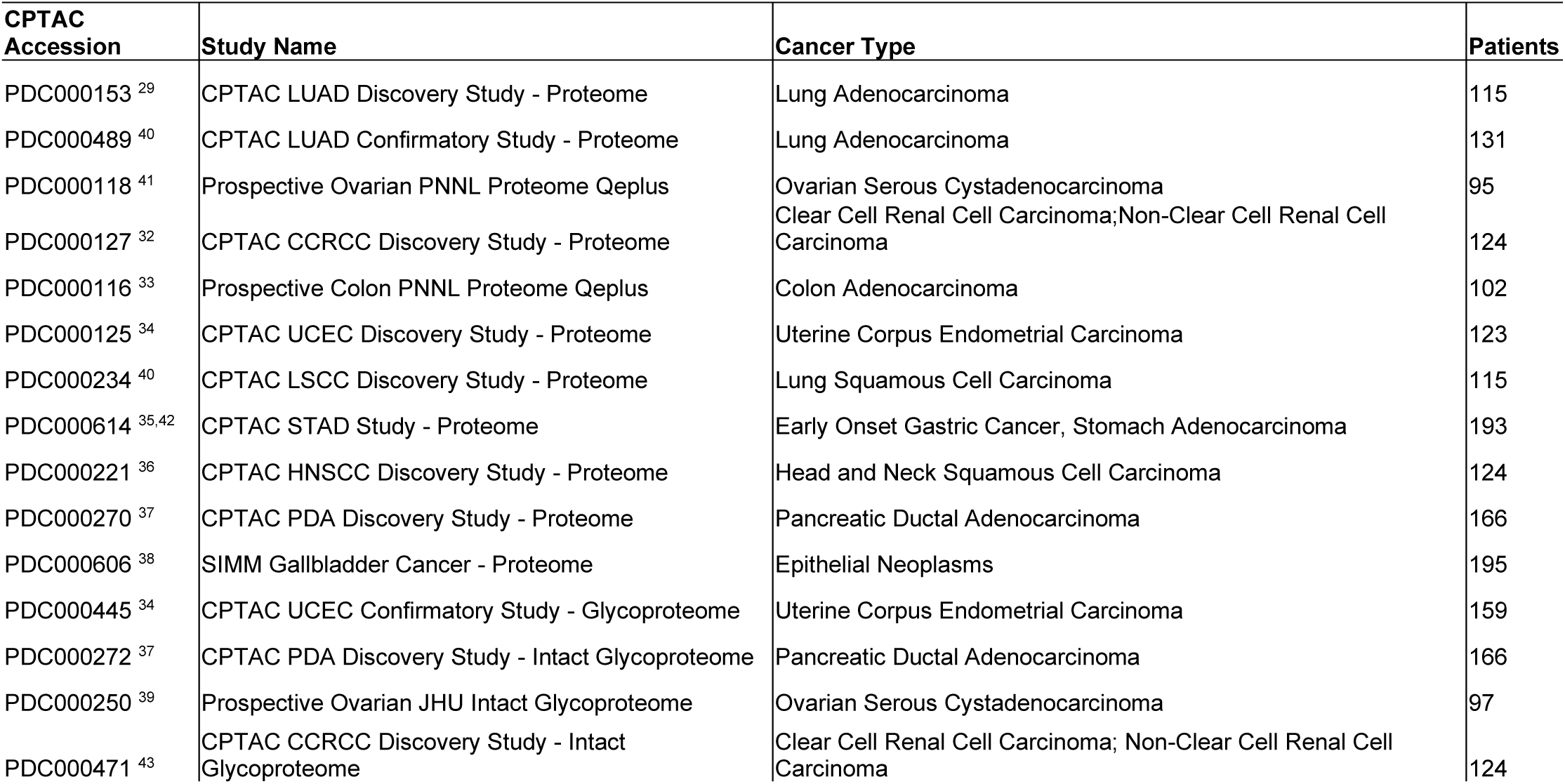
CPTAC Cellular and Membrane Proteomics Datasets. ^29,32–43^

**Supplementary Table 2.**
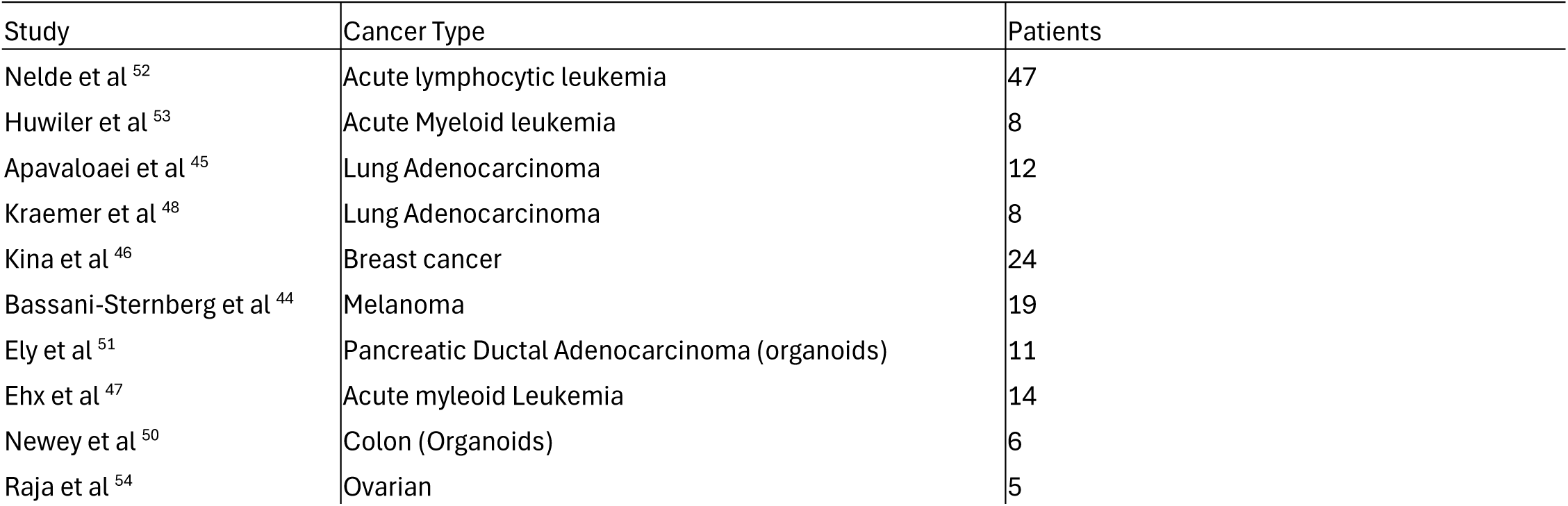
Publicly Available pMHC Datasets. ^44–54^

**Supplementary Fig. 2.**
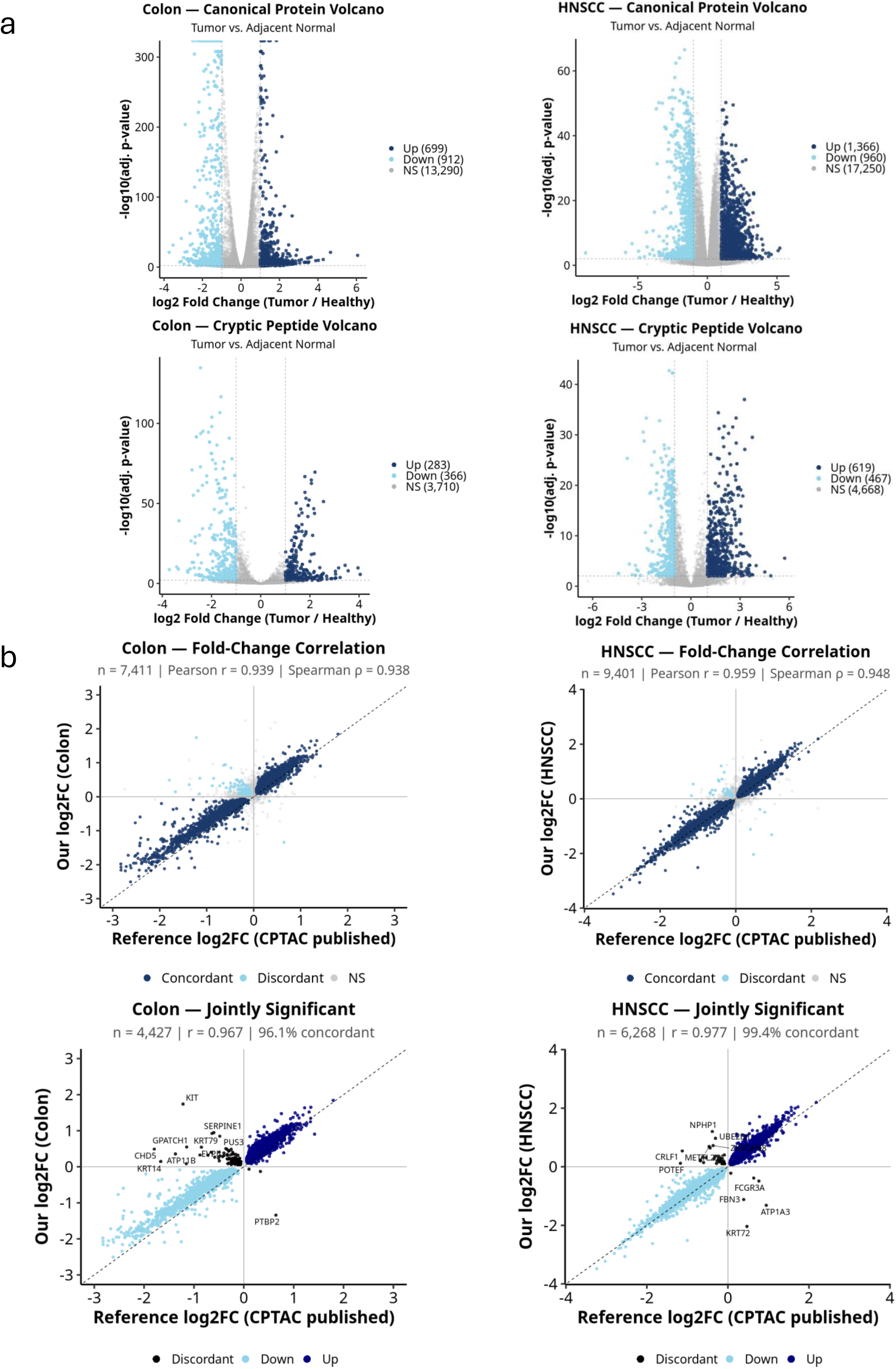
Discovery and initial characterization of a cryptic human proteome. **(a)** DarkRNA-encoded cryptic peptidome landscape across CPTAC cellular proteomic cohorts. Paired volcano plots of canonical protein (top row) and cryptic peptide (bottom row) differential abundance for Colon and HNSCC (Tumor vs. Adjacent Normal; adj. p < 0.01, |log2FC| >= 1). Colon canonical: Up (699), Down (912), NS (13,290); HNSCC canonical: Up (1,366), Down (960), NS (17,250). Colon cryptic: Up (283), Down (366), NS (3,710); HNSCC cryptic: Up (619), Down (467), NS (4,668). **(b)** Per-cohort cross-validation of canonical protein log2FC vs. published CPTAC reference for Colon and HNSCC. Top row: full fold-change correlations (Colon: n = 7,411, Pearson r = 0.939, Spearman r = 0.938; HNSCC: n = 9,401, Pearson r = 0.959, Spearman r = 0.948); bottom row: jointly significant subset with discordant outliers labeled (Colon: n = 4,427, r = 0.967, 96.1% concordant; HNSCC: n = 6,268, r = 0.977, 99.4% concordant).

**Supplementary Fig. 3.**
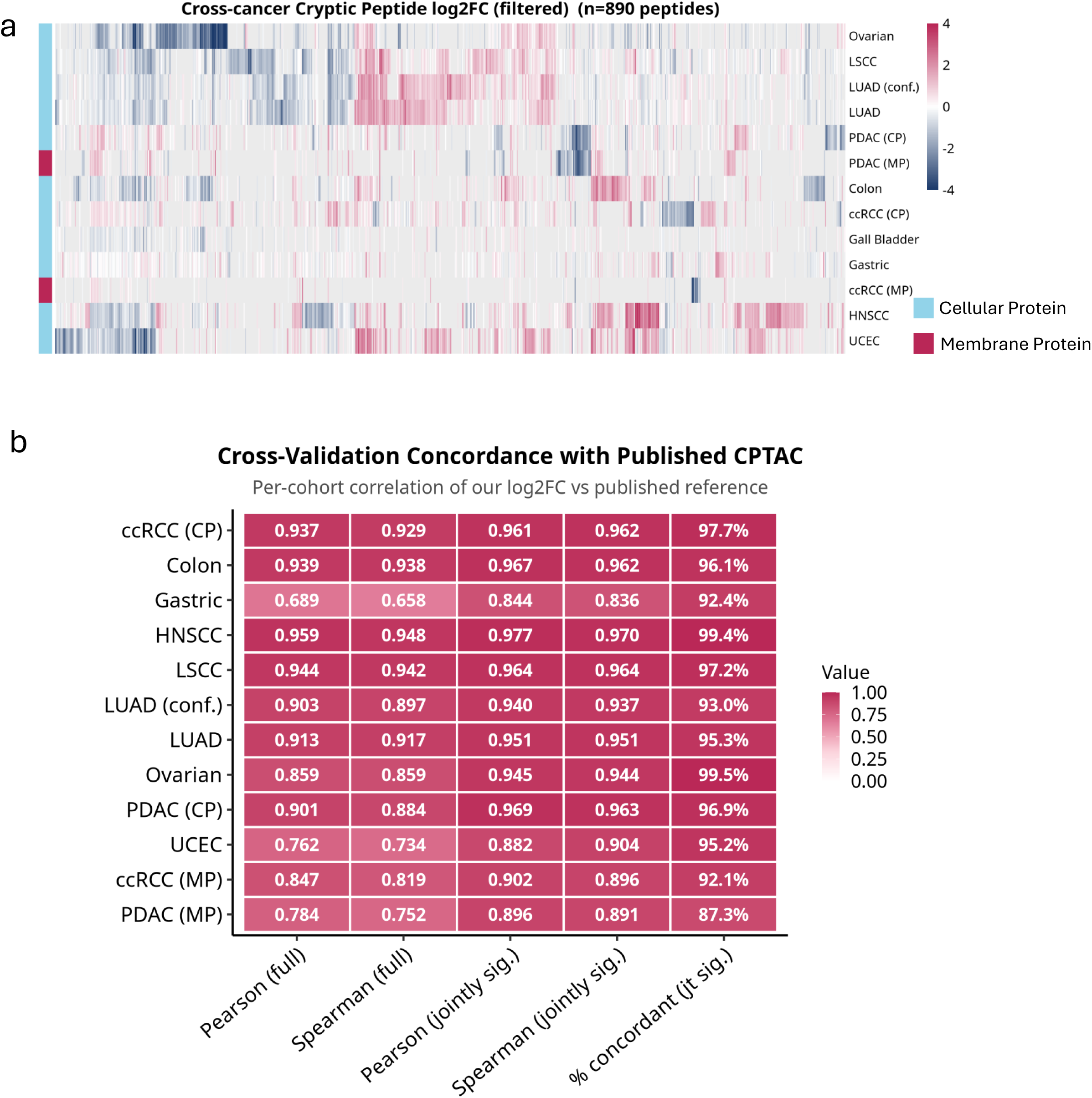
Initial characterization of a cryptic human proteome. **(a)** Cross-cancer cryptic peptide log2FC heatmap (filtered; n = 890 peptides; adj. p < 0.01, |log2FC| >= 1). Cohorts: Ovarian, LSCC, LUAD (conf.), LUAD, PDAC (CP), PDAC (MP), Colon, ccRCC (CP), Gallbladder, Gastric, ccRCC (MP), HNSCC, UCEC. Colored by assay: Cellular Protein (blue), Membrane Protein (pink). Rows hierarchically clustered; columns ordered by cohort. **(b)** Cross-validation concordance with published CPTAC. Per-cohort correlation of log2FC vs. published reference across 12 cohorts: ccRCC CP (Pearson 0.937, Spearman 0.929, jointly sig. Pearson 0.961, Spearman 0.962, 97.7% concordant), Colon (0.939, 0.938, 0.967, 0.962, 96.1%), Gastric (0.689, 0.658, 0.844, 0.836, 92.4%), HNSCC (0.959, 0.948, 0.977, 0.970, 99.4%), LSCC (0.944, 0.942, 0.964, 0.964, 97.2%), LUAD conf. (0.903, 0.897, 0.940, 0.937, 93.0%), LUAD (0.913, 0.917, 0.951, 0.951, 95.3%), Ovarian (0.859, 0.859, 0.945, 0.944, 99.5%), PDAC CP (0.901, 0.884, 0.969, 0.963, 96.9%), UCEC (0.762, 0.734, 0.882, 0.904, 95.2%), ccRCC MP (0.847, 0.819, 0.902, 0.896, 92.1%), PDAC MP (0.784, 0.752, 0.896, 0.891, 87.3%).

**Supplementary Fig. 4.**
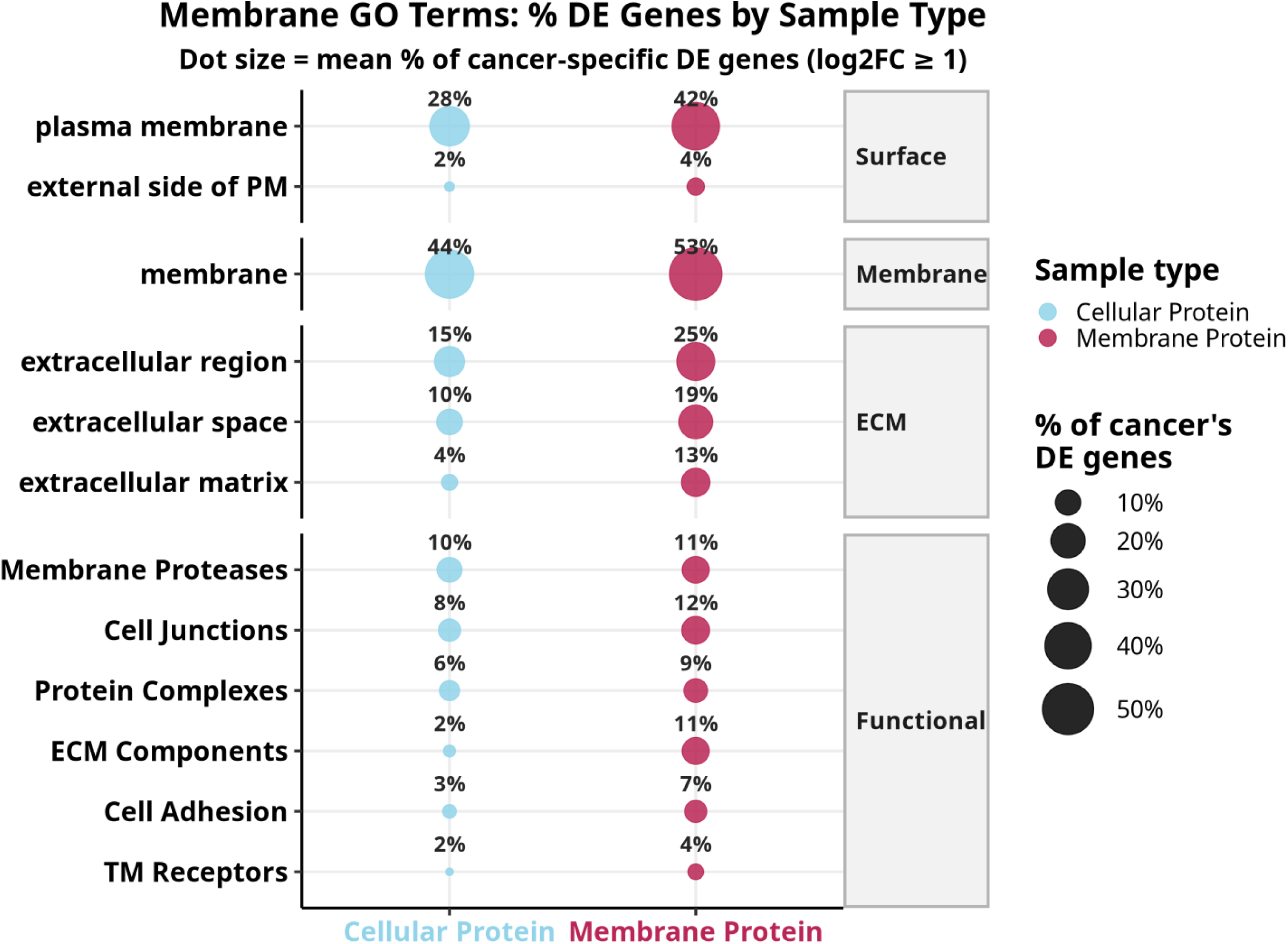
Selection of actionable membrane targets from the cryptic cancer proteome. Membrane GO Terms: mean percentage of cancer-associated DE genes (log2FC >= 1) by sample type. Dot size encodes mean percentage across cancers; Cellular Protein (light blue), Membrane Protein (pink). Surface terms: plasma membrane (28% Cellular, 42% Membrane), external side of PM (2%, 4%). Membrane: 44% Cellular, 53% Membrane. ECM terms: extracellular region (15%, 25%), extracellular space (10%, 19%), extracellular matrix (4%, 13%). Functional terms: Membrane Proteases (10%, 11%), Cell Junctions (8%, 12%), Protein Complexes (6%, 9%), ECM Components (2%, 11%), Cell Adhesion (3%, 7%), TM Receptors (2%, 4%). Membrane Protein consistently enriches surface-accessible and ECM-associated categories at higher fractions than Cellular Protein, supporting the use of membrane proteomics for identifying cryptic targets with extracellular accessibility.

**Supplementary Fig. 5.**
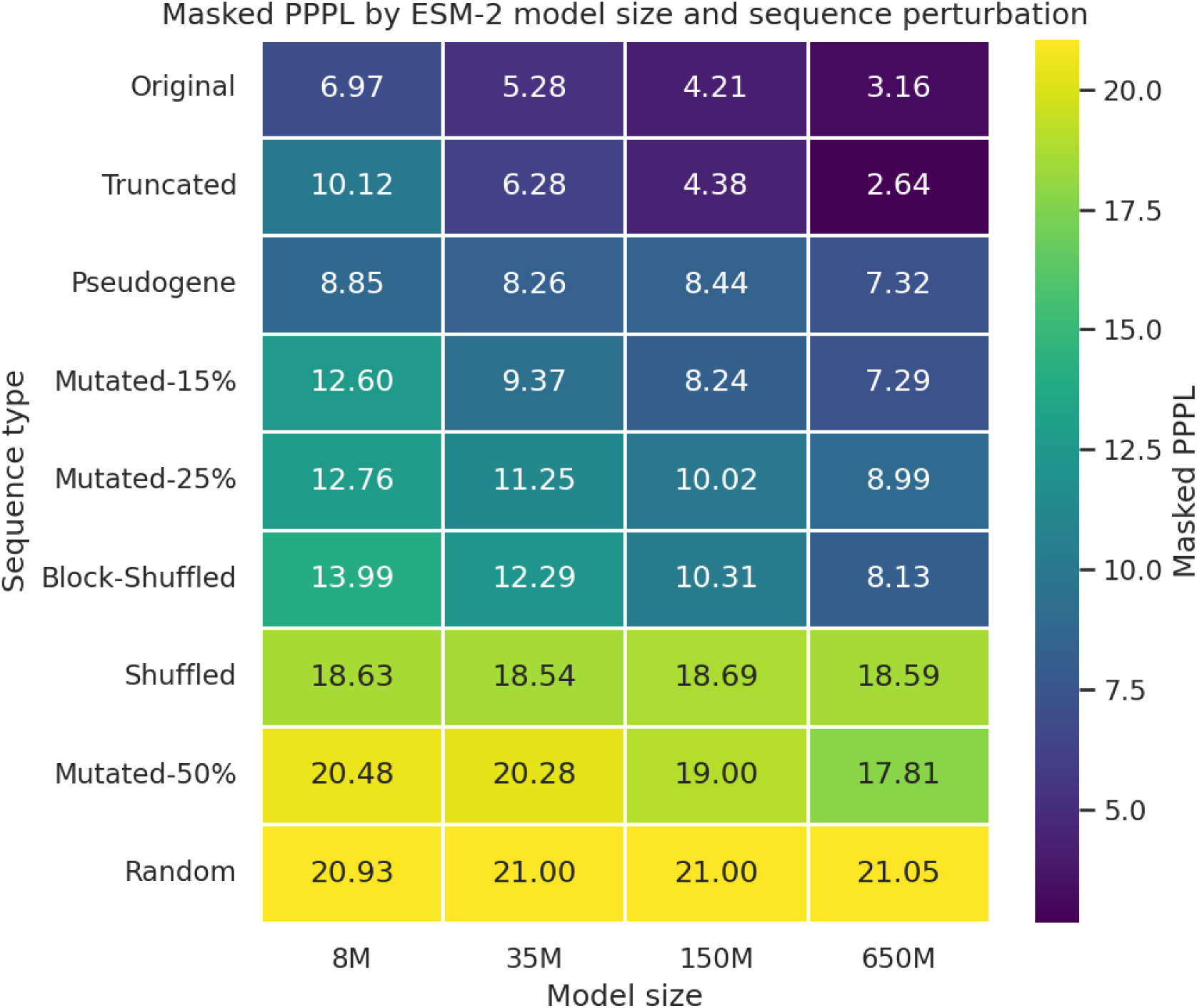
Pseudo-perplexity tracks perturbation severity across ESM-2 model sizes. Standard masked pseudo-perplexity (PPPL) for the canonical reference protein CP3A4_HUMAN and a graded panel of synthetic perturbations (an N-terminal truncation, a block shuffle, point mutations at 15%, 25%, and 50%, a fully shuffled sequence, a uniformly random sequence, and a same-length pseudogene-derived translation) evaluated at four ESM-2 backbones (8M, 35M, 150M, and 650M parameters). Rows are sorted by mean PPPL; lower values indicate sequences closer to natural protein space. Mean PPPL increases monotonically with perturbation severity, from 4.9 for the unperturbed sequence to 21.0 for the uniformly random control, with the pseudogene-derived translation at an intermediate value of 8.2, and the ordering is stable across model sizes.

**Supplementary Fig. 6.**
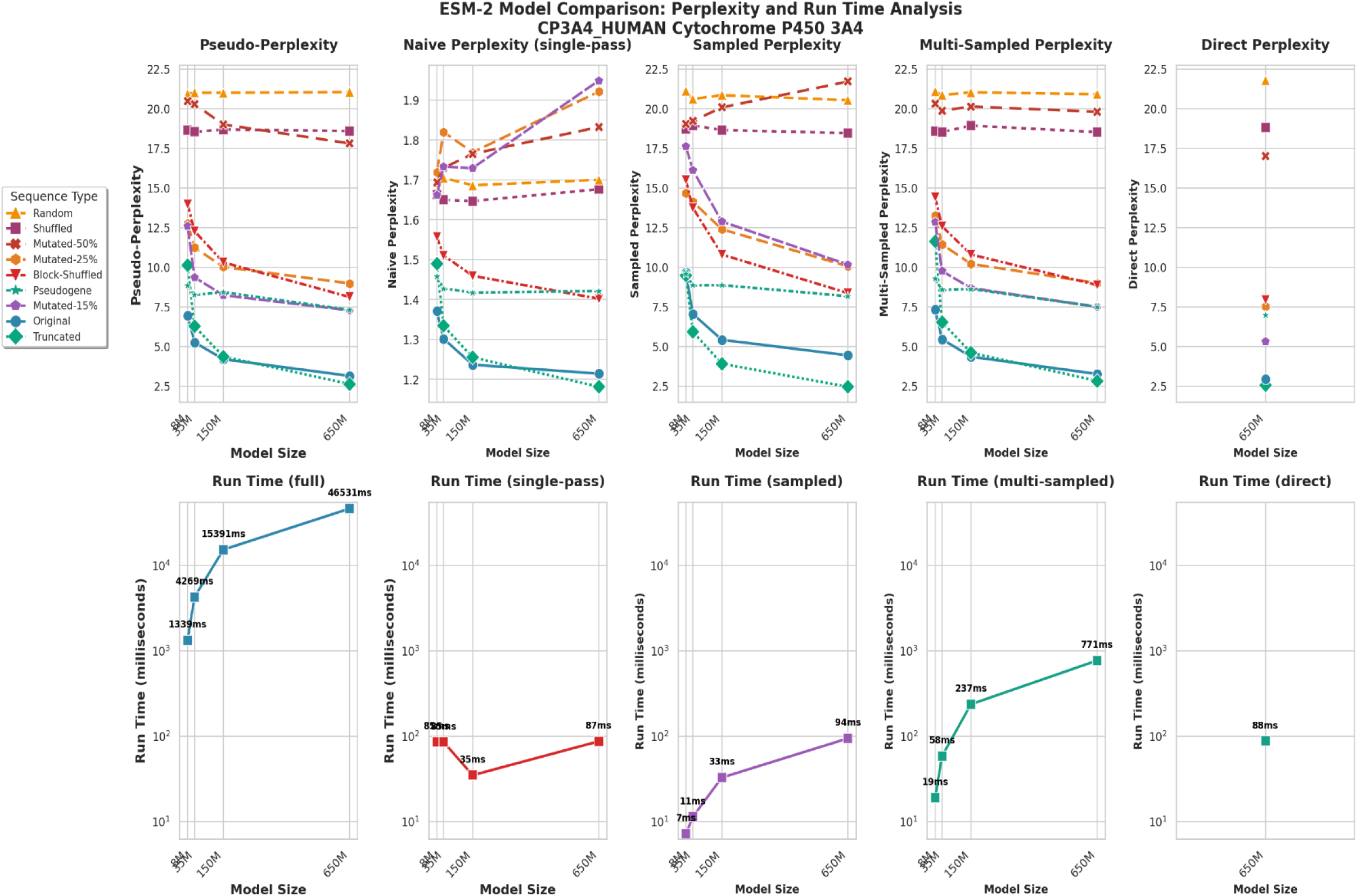
Speed–fidelity comparison of five PPPL protocols. Five PPPL protocols (standard masked, naive, sampled, multi-sampled, and direct) were evaluated on the nine-class sequence-perturbation benchmark across the four ESM-2 model sizes. Top row, PPPL versus model size (one line per sequence class); bottom row, mean inference run time (logarithmic scale). Standard masked PPPL separates the sequence classes cleanly with an ordering stable across model sizes; naive PPPL collapses to a narrow, non-discriminative band; sampled and multi-sampled PPPL preserve the masked ordering with progressively lower variance; and direct PPPL reproduces the masked ordering at single-pass cost.

**Supplementary Fig. 7.**
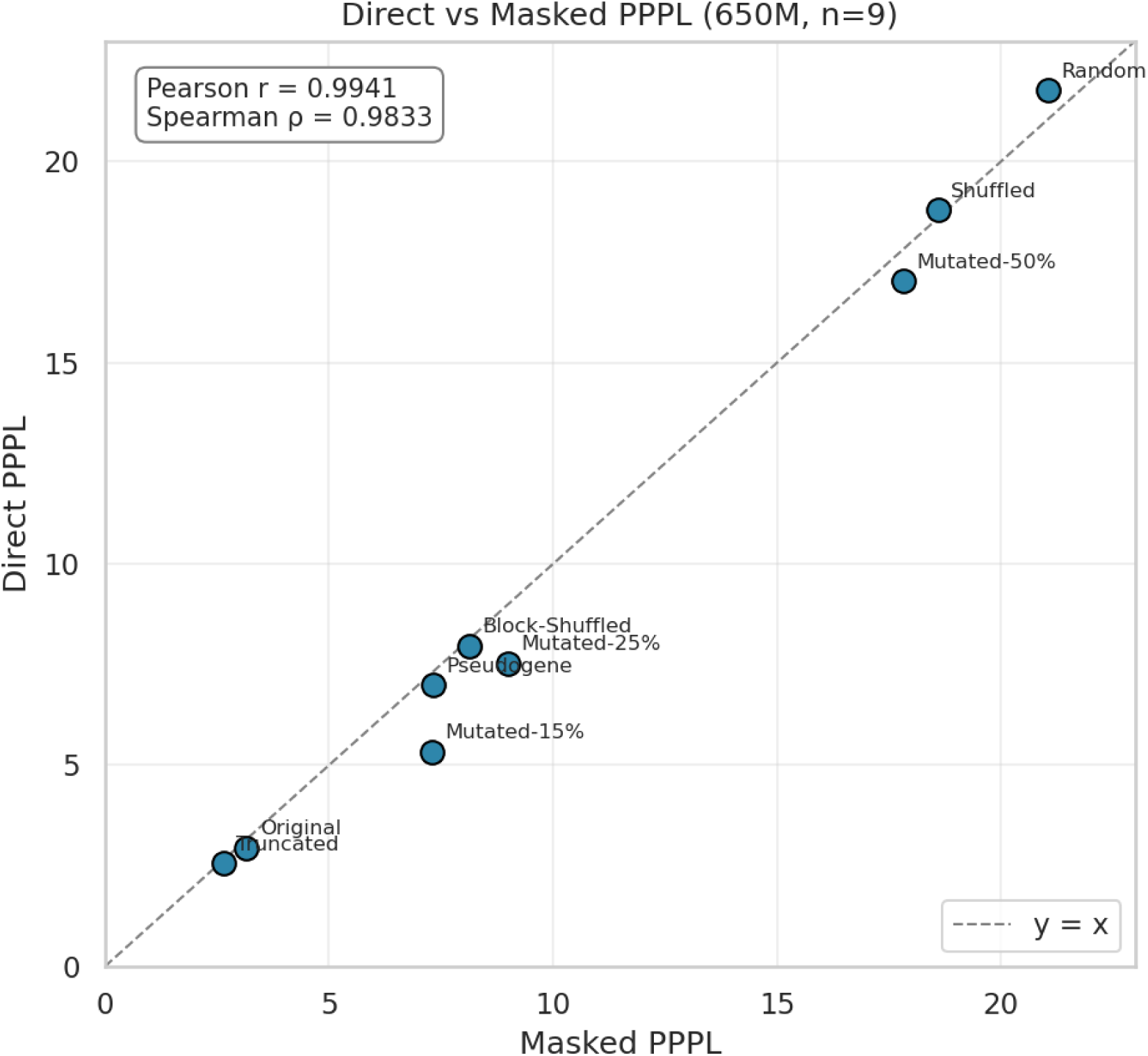
Direct PPPL reproduces standard masked PPPL. Direct versus standard masked PPPL across the nine sequence classes at the 650M ESM-2 backbone; the dashed line is the y = x identity. Direct and masked PPPL agree closely across an approximately seven-fold range of PPPL (Pearson r = 0.994, Spearman ρ = 0.983), with the largest discrepancies confined to the mid-perturbation regime.

**Supplementary Fig. 8.**
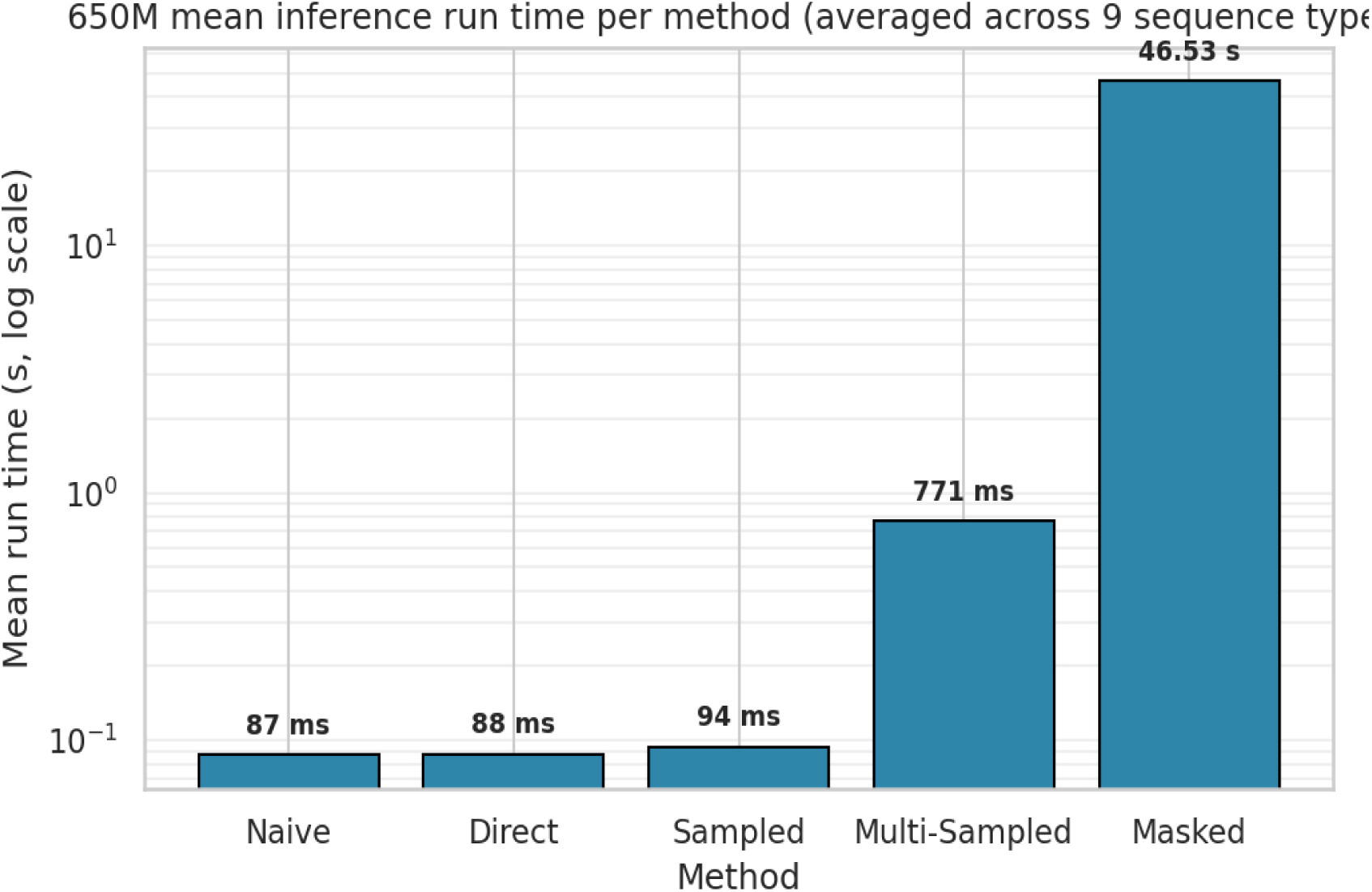
Direct PPPL is orders of magnitude faster than masked PPPL. Mean inference run time per PPPL protocol at the 650M ESM-2 backbone, averaged across the nine sequence classes (logarithmic y-axis). Direct PPPL averages 88 ms per sequence versus 46.5 s for standard masked PPPL (an approximately 530-fold reduction) while preserving the fidelity shown in Supplementary Fig. 7.

**Supplementary Fig. 9.**
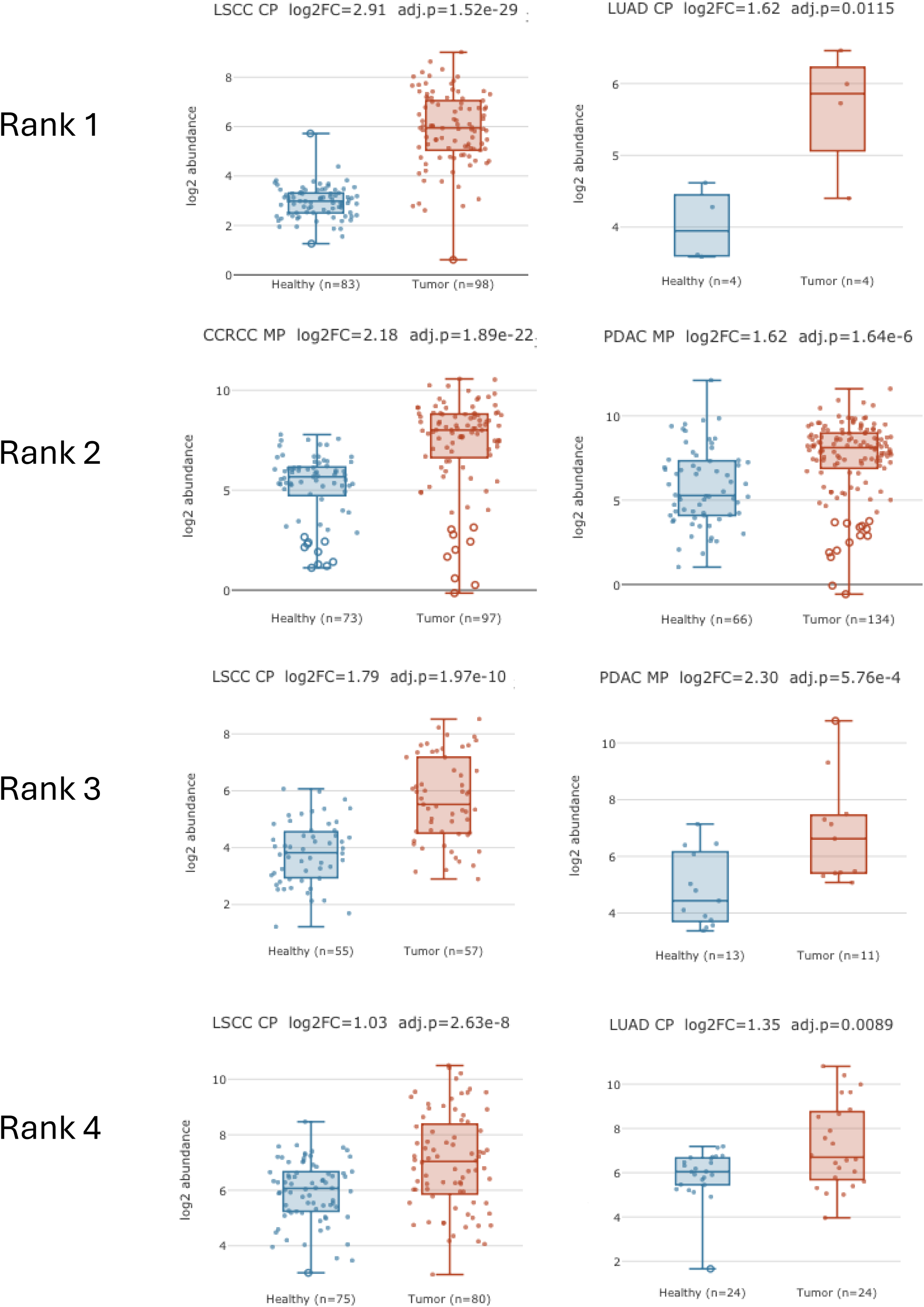
Differential expression of peptides mapping to top ranked cryptic membrane proteins. Tumor vs. healthy log2 abundance boxplots for the top four ranked membrane target cryptic ORFs from Fig. 4a, shown across their most differentially expressed CPTAC cohorts. Rank 1 (RD-8886583): Lung Squamous Cell Carcinoma (LSCC) cellular proteomics log2FC = 2.91, adj. p = 1.52e-29 (Healthy n = 83, Tumor n = 98); Lung Adenocarcinoma (LUAD) cellular proteomics log2FC = 1.62, adj. p = 0.0115 (Healthy n = 4, Tumor n = 4); Clear Cell Renal Cell Carcinoma (ccRCC) membrane proteomics log2FC = 2.18, adj. p = 1.89e-22; Pancreatic Ductal Adenocarcinoma (PDAC) membrane proteomics log2FC = 1.62, adj. p = 1.64e-6. Rank 2 (RD-5336591): LSCC cellular proteomics (Healthy n = 73, Tumor n = 97); PDAC membrane proteomics (Healthy n = 66, Tumor n = 134). Rank 3 (RD-0376778): LSCC cellular proteomics log2FC = 1.79, adj. p = 1.97e-10 (Healthy n = 55, Tumor n = 57); PDAC membrane proteomics log2FC = 2.30, adj. p = 5.76e-4 (Healthy n = 13, Tumor n = 11). Rank 4 (RD-8653598): LSCC cellular proteomics log2FC = 1.03, adj. p = 2.63e-8 (Healthy n = 75, Tumor n = 80); LUAD cellular proteomics log2FC = 1.35, adj. p = 0.0089 (Healthy n = 24, Tumor n = 24). Blue = healthy, red = tumor.

**Supplementary Fig. 10.**
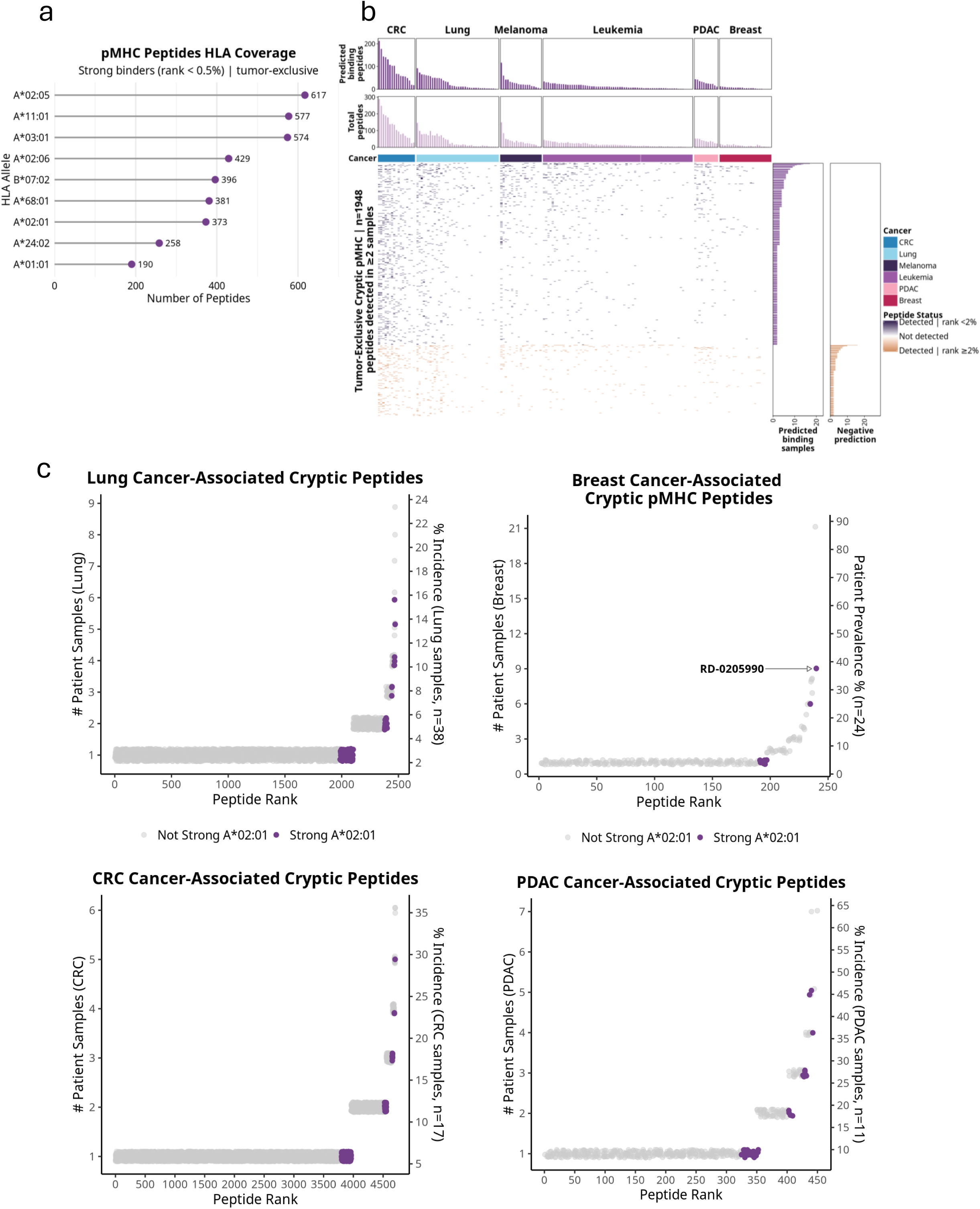
Selection of actionable targets from the cryptic cancer immunopeptidome - extended data. **(a)** pMHC Peptides HLA Coverage: tumor-exclusive cryptic strong binders (MHCflurry rank <0.5%) per HLA allele. A*02:05 (617), A*11:01 (577), A*03:01 (574), A*02:06 (429), B*07:02 (396), A*68:01 (381), A*02:01 (373), A*24:02 (258), A*01:01 (190). **(b)** Per-sample detection matrix of tumor-exclusive cryptic pMHC peptides detected in >=2 samples (n = 1,948 peptides) across CRC, Lung, Melanoma, Leukemia, PDAC, and Breast. Purple = detected, rank <2%; orange = detected, rank >=2%; grey = not detected. **(c)** Ranked patient prevalence scatter plots for tumor-exclusive HLA-A*02:01 strong binder cryptic pMHC peptides in Lung (n = 38), Breast (n = 24), CRC (n = 17), and PDAC (n = 11). Purple = strong A*02:01 binder; grey = not strong. RD-0205990 annotated.

**Supplementary Fig. 11.**
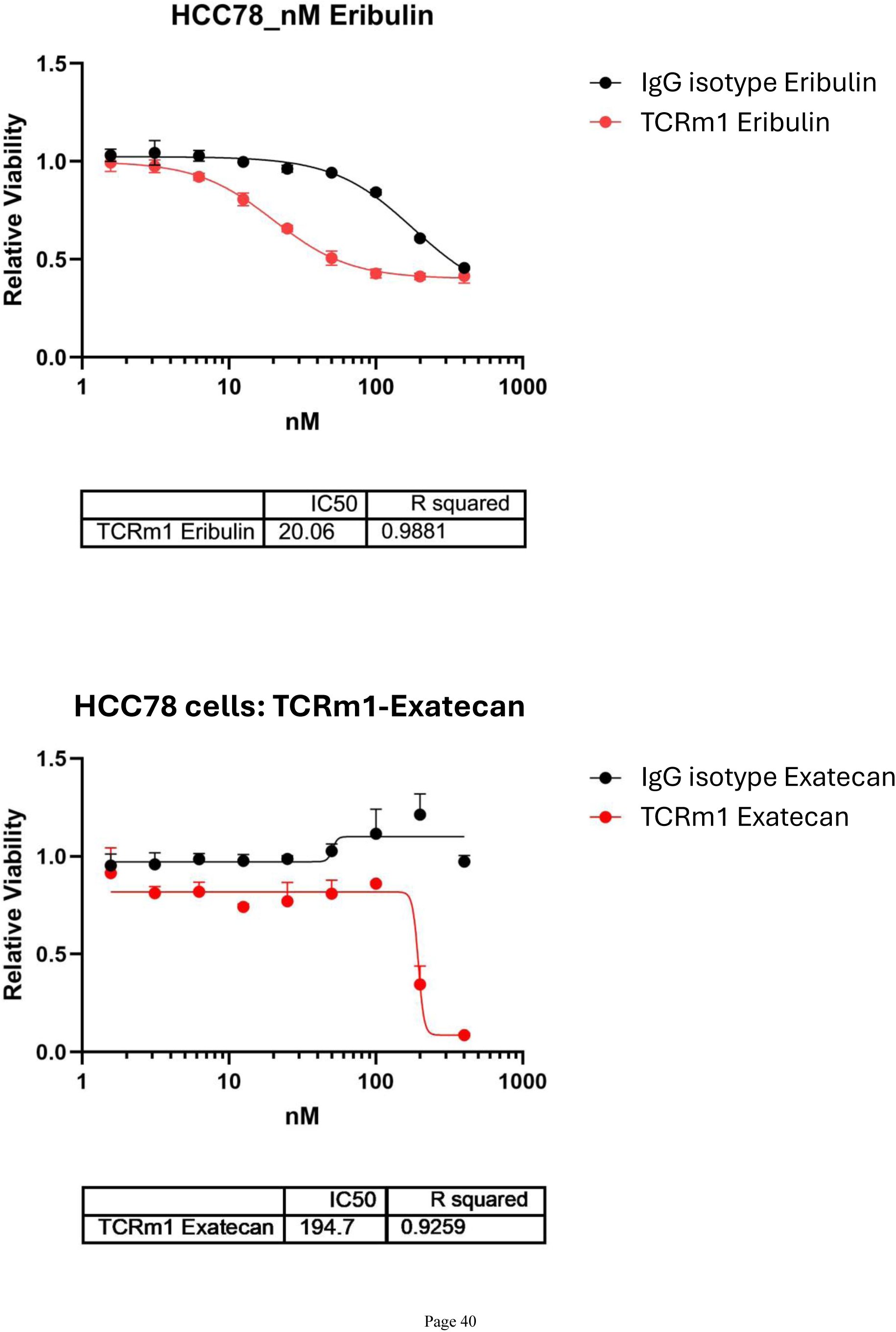
Demonstration of *in vitro* cancer cell killing by TCRm1 formatted as ADCs. (Top) TCRm1 was reduced with TCEP, and Eribulin was conjugated via maleimide to free sulfhydryl residues (cysteines) through a lysosome-cleavable linker, to arrive at an average drug to antibody ratio (DAR) of ∼6 (red). The same reaction conditions were used for linker/payload (L/P) derivatization of an anti-beta galactosidase human IgG1 antibody (IgG isotype control-Eribulin (black)). HCC78 cells were plated (3e3 cells/well) in 96 well plates and treated with a serial dilution of antibody (400, 200, 100, 50, 25, 12.5, 6.25, 3.12, and 1.56nM). Plates were incubated for 84h before cell viability determination was performed with CelTiter-Glo 2.0. Exposure of cells to 50nM TCRm1-Eribulin achieved a relative viability score of 0.5 in this assay (calculated IC_50_ ∼20nM), while the IgG isotype control-Eribulin conjugate approached this value only at the highest concentrations tested (200 and 400nM). **(Bottom)** TCRm1 was conjugated to a maleimide-functionalized, lysosome-cleavable linker to Exatecan under the same reaction conditions used for derivatization with Eribulin, with slightly lower DAR achieved (∼4.5 (red)). The Exatecan L/P was also conjugated to the same anti-beta galactosidase human IgG1 antibody (IgG isotype control-Exatecan (black)). HCC78 cells were plated (3e3 cells/well) in 96 well plates and treated with a serial dilution of antibody (400, 200, 100, 50, 25, 12.5, 6.25, 3.12, and 1.56nm). Plates were incubated for 84h before cell viability determination was performed with CelTiter-Glo 2.0. Cells exposed to 200nM and 400nM TCRm1-Exatecan had significantly reduced viability (∼30% and ∼5% respectively) relative to all concentrations of the IgG isotype control-Exatecan tested in this assay. IC_50_ = 194.7nM.

## Methods

### Tissue Sample Preparation

Tumor and healthy tissue samples were obtained from Crown Bio. Samples were pulverized using the cryoPREP Manual Dry Pulverizer (Covaris, CP-01) and aliquoted for nucleic acid and protein extractions.

### Nucleic Acid Extractions

RNA and DNA were extracted from tissue samples using TRIzol Reagent (ThermoFisher Scientific, 15596026) according to the manufacturer’s protocols for RNA and DNA isolation. Total RNA was treated with DNAseI (NEB, M0303L) for 30 minutes at 37°C to remove genomic DNA and cleaned up with RNAClean XP RNA and cDNA Cleanup Reagent (Beckman Coulter, A63987). RNA and DNA were quantified by nanodrop.

### HLA Typing

Tissues were HLA typed for HLA-A, HLA-B, and HLA-C using primers described previously (Stockton et al., 2020). 30 ng gDNA was combined with 1X LongAmp Taq MM (NEB, M0287S) and 400 nM of each primer (IDT) in 25 uL and cycled according to manufacturer protocols for 30 cycles with an annealing temperature of 62C and a four minute 30 second extension time. Reactions were purified with AMPure XP beads (Beckman Coulter, A63881), prepared for sequencing using the Native Barcoding 24 V14 protocol (Oxford Nanopore Technologies), and sequenced on MinION R10 flow cells with barcode trimming enabled. Samples were haplotyped using the HLA-LA workflow (https://github.com/DiltheyLab/HLA-LA).

### pMHC Sample Preparation

HLA-peptide complexes were isolated from tumor tissue and cell lines by immunoaffinity purification. Briefly, patient tissue or cell pellets (∼200 mg tissue or ∼5×108 cells) were lysed in pMHC lysis buffer (0.25% sodium deoxycholate, 0.2 mM iodoacetamide, 1 mM EDTA, 1:200 protease inhibitor cocktail, 1 mM PMSF in PBS) for 15 mins rotating at 4C. Lysates were clarified by centrifugation at 20,000 × g for 20 min at 4°C. Pan-HLA class I antibody (Biolegend, San Diego; Catalog Number: 311402) was used for immunoprecipitation. W6/32 antibody was coupled to protein A agarose beads for an hour rotating at RT. Beads were then washed with 10x PBS before incubation with 10mM DMP for 30 mins, twice. Beads were then washed with 5 ml of PBS, 5 ml 100 mM Tris (pH 8) before incubating with 5% ethanolamine for 2 hrs at RT. Beads were then washed with 5 ml of PBS. Beads were then added to cleared lysates and rotated overnight at 4C. Beads were washed with 5 ml of pMHC lysis buffer, 5 ml PBS, 5 ml 10 mM Tris (pH 8). Peptides were eluted from pMHC complexes using 3x 200 ul 0.1% TFA. Samples were then desalted using C18 and then vacuum dried for LC-MS/MS analysis.

## LC-MS/MS Analysis

### Orbitrap Ascend

HLA peptides were analyzed using either a Thermo Orbitrap Exploris 480 or Thermo Orbitrap Ascend Tribrid mass spectrometer (Thermo Scientific) coupled with a Vanquish Neo UHPLC system (Thermo Scientific) and equipped with a FAIMS Pro Duo interface or with the FAIMS interface omitted. FAIMS was omitted only for a few samples included in the analysis in Figure 1 and 2 during the early validation of the KP/Ab Strep model. Samples were resuspended in 2% ACN +0.1% FA and directly loaded onto a precolumn (2 cm x μm ID packed with C18 reversed-phase resin, 5μm, 100 Å) in solvent A (2% ACN, 0.1% FA). Trapped peptides were eluted onto the analytical capillary chromatography column with an integrated electrospray tip (C18, 75 μm ID × 25 cm, 2 μm, 100 Å, ThermoScientific). Peptides were eluted over 105 minutes with the following: gradient 2–3% solvent B (90% ACN, 0.1% FA) for 10 min, 3–5% solvent B for 2 minutes, 5–23% solvent B for 63 minutes, 23-40% solvent B for 22 minutes, 40-90% solvent B for 3 minutes, hold for 5 minutes and followed by column equilibration and wash.

Data dependent acquisition was performed in positive ion mode at a spray voltage of 2.0 kV and heated capillary temperature, 275°C. Full scan mass spectra (300–1,000 m/z, 120,000 resolution for MHC-I and 300-1,200 m/z, 120,000 resolution for MHC-II) were detected in the Orbitrap analyzer after accumulation of 2.5 × 106 ions (normalized automatic gain control (AGC) target of 250%) and 250 ms maximum injection time (IT). For every full scan, MS2 spectra were collected during a 1.5 s cycle time for FAIMS compensation voltages at -45 and -65 or 3 s cycle time without FAIMS. Precursor ions were isolated with a 2.0 m/z isolation window and accumulation time was set to auto with 250 ms maximum accumulation time. Ions were fragmented with higher energy disassociation collision (HCD) with 30% collision energy (CI) at a resolution of 30,000. Charge states <2 and >4 were excluded for MHC-I and <2 and >6 were excluded for MHC-II, and dynamic exclusion was set to 15 s after n = 1 observation.

### Orbitrap Astral

Mass spectrometric data were collected on a Orbitrap Astral instrument coupled to a Vanquish Neo UHPLC. Peptides were separated using a 75-min gradient of 5 to 29% acetonitrile in 0.1% formic acid with a flow rate of 350 nL/min. The spray voltage was set at 2700 V. The scan sequence began with an Orbitrap MS1 spectrum with the following parameters: resolution 120000, scan range 350-1350 Th, automatic gain control (AGC) target 1000000, and maximum injection time 50 ms. MS2 spectra were acquired in a data-dependent manner with a scan cycle Top30 using the following parameters: resolution “Astral”, AGC target 20000, maximum injection time 15 ms, isolation window 1.2 Th, normalized collision energy (NCE) 27.00%, and centroid spectrum data type. Dynamic exclusion was set to automatic. The FAIMS compensation voltages (CV) were -25, -35, -45, -55 and-65 V.

### CPTAC Cellular and Membrane Proteomics

Raw TMT mass spectrometry files from the Clinical Proteomic Tumor Analysis Consortium (CPTAC) were downloaded from the CPTAC Data Portal (https://proteomics.cancer.gov/data-portal). Cellular TMT proteomics datasets were analyzed for colorectal carcinoma (CRC; PDC000116), lung squamous cell carcinoma (LSCC; PDC000234), and lung adenocarcinoma (LUAD; PDC000153) cohorts. Glycoproteome enriched TMT datasets were analyzed for pancreatic ductal adenocarcinoma (PDAC; PDC000270), ovarian cancer (OV; PDC000250), uterine corpus endometrial carcinoma (UCEC), and clear cell renal cell carcinoma (ccRCC) cohorts. Downloaded raw spectra were searched using Comet against a custom protein sequence database comprising the canonical human proteome and RyboCypher-derived cryptic protein predictions, prior to downstream quantitative processing.

### Comet Database Search Parameters

All raw mass spectrometry spectra, including in-house or publicly available immunopeptidomics, CPTAC proteome and glycoproteome proteomics datasets were searched using Comet (2026.01 rev.1) against a custom protein sequence database. Both CPTAC and immunopeptidomics datasets are listed with references below in Supp Table 1 and 2. The database comprised the Ensembl human reference proteome (103), RyboCypher-derived cryptic protein predictions and nuORF database entries^14^. For immunopeptidomics searches, no enzyme specificity was applied to accommodate the non-tryptic nature of HLA ligands; for CPTAC TMT datasets, trypsin was specified as the digestion enzyme with up to 2 missed cleavages permitted. Search parameters for pMHC data included: precursor mass tolerance of 20 ppm, fragment ion tolerance of 0.02 Da, variable modifications of oxidation on methionine (+15.995 Da), and peptide length restricted to 8–16 amino acids. To minimize the detrimental effects of search space using such large databases with no enzyme searches, our original lung and colon datasets used for Rybocypher sequencing were searched against the full proteomics database as a first pass. Peptides that were identified using very lenient false positives rates of 25% were then used to create a second database of ∼600,000 sequences to perform a secondary database search where a 1% false positive rate was used for peptide assignments. For TMT datasets, static modifications of +229.163 Da for TMT11 plex labeling on lysine and on the N-terminus, carbamidomethylation of cysteine (+57.021 Da), with the same precursor and fragment tolerances applied. Like immunopeptidomics datasets, TMT data was filtered to a 1% false positive rate. False positive rates were determined using a target-decoy approach^55^ using a linear discriminant score.

## CPTAC MSstatsTMT

### Canonical Protein Differential Expression

CPTAC TMT PSM-level quantification data were processed using the MSstatsTMT Bioconductor package^24^. PSMs were filtered at a 1% LDA Q-value threshold with contaminant proteins excluded prior to analysis. Protein-level abundances were estimated from PSM-level signal-to-noise ratios using Tukey’s median polish summarization, with global and reference-channel (bridge) normalization applied to correct for loading differences and between-plex batch effects. Differential expression between tumor and matched adjacent-normal samples was assessed using a linear mixed-effects model with empirical Bayes variance moderation, accounting for biological replicate and TMT plex as random effects. Multiple testing correction was applied using the Benjamini– Hochberg (BH) procedure; proteins with an adjusted p-value < 0.05 were considered significant.

### DarkRNA-Derived Peptide Identification and Differential Expression

To identify and quantify peptides derived from DarkRNA sequences within CPTAC TMT datasets, PSM-level data were searched against a custom DarkRNA protein sequence database alongside the canonical proteome. Candidate DarkRNA peptides were validated by tryptic digest mapping and a sequence-specificity filter requiring confirmed mismatches to the canonical human proteome, ensuring the identified peptides could not be explained by known protein sequences. Protein summarization and normalization were performed on the complete PSM dataset prior to subsetting, preserving stable normalization across the full dynamic range of the experiment. Validated DarkRNA peptides were then extracted and differential expression between tumor and adjacent-normal was assessed using the same MSstatsTMT linear mixed-effects framework and BH correction as for canonical proteins.

### Validation against Published CPTAC Reference Data

Differential expression results were benchmarked against independently published CPTAC proteome data retrieved from LinkedOmics (http://linkedomics.org). Reference differential expression was computed using a statistical test matched to each dataset’s structure: a one-sample t-test for pre-computed per-patient log2 fold-change matrices (CRC), a paired t-test for cohorts with matched tumor/normal abundance matrices (LSCC, LUAD), and Welch’s t-test for cohorts with unpaired samples (OV, PDAC). For glycoproteomics reference datasets (OV, PDAC), site-level abundances were aggregated to gene level by median prior to comparison. Agreement was evaluated by Pearson and Spearman correlation of gene-level log2 fold-changes, directional concordance, and Bland–Altman analysis. BH correction was applied to all reference p-values^56^.

### Sequence divergence scoring of cryptic peptides from the canonical proteome

Peptides identified by mass spectrometry were scored for sequence similarity to the canonical human proteome using Levenshtein (edit) distance, which quantifies the minimum number of substitutions, insertions, and deletions required to align a query sequence to any substring in the reference. Scoring was performed using the edlib Python library (v1.3.9) in infix (“HW”) alignment mode against a concatenated target comprising all canonical human reference proteins (Gencode v42, hg38), with sentinel characters between proteins to prevent cross-protein alignments. Peptides with an edit distance of zero, i.e. exact substring matches to any canonical protein, were excluded. The edit distance was capped at 5; peptides exceeding this threshold were recorded as greater than 5.

### Gene Ontology enrichment of upregulated cryptic peptides

Each cryptic peptide was annotated with the gene symbol(s) of overlapping canonical loci identified during peptide-genome alignment. For each cancer cohort, gene symbols were extracted from significantly upregulated cryptic peptides (coreMismatchNumber >= 1, adjusted p < 0.01, log2FC >= 1), split, and deduplicated to produce a unique gene list. The statistical background was defined as all unique genes annotated to any Rybodyn-detected peptide (coreMismatchNumber >= 1) across all cohorts.

GO:BP enrichment was performed independently for each of the 11 Cellular-MS cohorts (HNSCC, LSCC, LUAD, Colorectal, PDAC, UCEC, Ovarian, ccRCC, Gastric, Gallbladder, and a LUAD confirmation cohort) and 4 Membrane-MS cohorts (PDAC, ccRCC, Ovarian, UCEC) using the gprofiler2 R package (gost() function, Homo sapiens, Benjamini-Hochberg FDR correction, q < 0.05)^57^. Fold enrichment was calculated as (intersection size / query size) / (term size / background size). Results were combined across cohorts and annotated by assay type (Cellular-MS or Membrane-MS) for visualization.

### HLA Binding Predictions

Predicted HLA class I binding affinities were computed for all candidate cryptic peptides using MHCflurry 2.0^58^. Peptides were scored against a panel of clinically prioritized HLA alleles selected based on global population prevalence, including HLA-A*02:01, HLA-A*02:05, HLA-A*02:06, and additional alleles as indicated. Both binding affinity (IC50, nM) and antigen presentation scores were generated using the MHCflurry presentation predictor, which integrates binding affinity, cleavage, and expression context. Peptides were classified as strong binders at a presentation percentile rank threshold of <0.5% and as weak binders at <2%, consistent with standard immunopeptidomics conventions. All predictions were run on peptides of length 8–11 amino acids. Allele-level results were integrated with immunopeptidomics detection data and patient HLA haplotyping to compute per-allele patient prevalence across each tumor cohort.

### Membrane Topology and Subcellular Localization Predictions

Transmembrane topology and subcellular localization of the top 10 cryptic protein candidates were assessed using two independent web server-based predictors. Transmembrane domain architecture was predicted using DeepTMHMM^27^ (https://dtu.biolib.com/DeepTMHMM), a deep learning-based topology predictor that classifies each residue as belonging to a transmembrane helix, intracellular loop, extracellular loop, or signal peptide. Full-length cryptic protein sequences were submitted individually, and topology annotations were recorded, including the number of predicted transmembrane segments, orientation (N-terminal inside vs. outside), and signal peptide probability. Subcellular localization was independently predicted using DeepLoc 2.0^28^ (https://services.healthtech.dtu.dk/services/DeepLoc-2.0), which provides multi-label localization probabilities across ten compartments including plasma membrane, nucleus, cytoplasm, extracellular space, and organelle membranes. Sequences were submitted using the default settings with the eukaryote organism parameter. Candidates were considered membrane-associated if DeepTMHMM predicted at least one transmembrane segment or a signal peptide, and DeepLoc assigned the highest localization probability to the plasma membrane or an endomembrane compartment. Concordance between the two predictors was used as an additional confidence filter for downstream prioritization.

### Proteasomal Processing Validation of Candidate ORFs

Candidate dark RNA ORFs containing immunopeptidomics-detected cryptic peptide sequences were evaluated for proteasomal processing compatibility using Pepsickle (v1.0.0; default model). Per-position cleavage probabilities were computed across the full ORF sequence; a site was considered cleaved if the predicted probability exceeded 0.6. ORFs were classified as supporting peptide liberation when a high-confidence cleavage event was present at the peptide C-terminus with no internal cleavage predicted within the peptide boundaries at the same threshold. Structural and functional domain annotations were generated with InterProScan (v5.78-109.0) using Phobius for transmembrane topology and cytoplasmic/extracellular region assignment, MobiDB-Lite for intrinsic disorder prediction, and PANTHER for functional domain classification. Regions sharing high similarity to canonical proteins are indicated as masked tracks in each topology panel.

## ESM-2 Pseudo-Perplexity Analysis

### ESM-2 Backbones

All sequence-plausibility analyses were performed using the ESM-2 family of protein language models^59,60^: the 8M-parameter (facebook/esm2_t6_8M_UR50D), 35M-parameter (facebook/esm2_t12_35M_UR50D), 150M-parameter (facebook/esm2_t30_150M_UR50D), and 650M-parameter (facebook/esm2_t33_650M_UR50D) checkpoints, loaded through the Hugging Face Transformers library. Inference was run in PyTorch with float32 precision on a single NVIDIA GPU; the 650M checkpoint was used for all cohort-scale analyses unless stated otherwise.

### Sequence Pseudo-Perplexity

For a protein sequence *x*_1_, …, *x*_*N*_ we computed sequence pseudo-perplexity (PPPL) as the exponential of the mean per-position token loss,

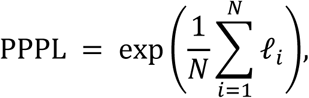

where the per-position loss ℓi was defined according to one of five protocols. Special tokens (CLS, EOS, padding) were excluded from the mean. For sequences exceeding the model’s maximum positional embedding, scores were computed over overlapping sliding windows of length max_length with a stride of max_length / 2, and per-position losses were averaged across overlaps before exponentiation.

Standard masked PPPL^60^ defined per-position loss as the negative log probability of the true residue given all other positions masked one at a time: ℓ_*l*_ = *log p_2θ_*(*x*_*l*_ | *x*_∖*l*_). For efficiency, mask positions were batched diagonally so that each forward pass scored a chunk of consecutive positions simultaneously.

Naive PPPL used the same expression but conditioned on the full unmasked sequence, ℓ_*l*_ = *log p_2θ_*(*x*_*l*_ | *x*), evaluated in a single forward pass. Because the model observes the target residue in its own input, naive PPPL exhibits substantial information leakage and was retained only as a diagnostic baseline.

Sampled PPPL scored the model on a random 15% subset of positions in a single forward pass, following the position-selection rule of the ESM-2 masked-language-modeling pretraining objective. A subset *S* ⊆ {1, …, *N*} of valid positions was drawn with selection probability *p_sel_* = 0.15. The implementation supports the full pretraining-style replacement scheme in which selected positions are independently masked, replaced with a uniformly random standard amino acid, or left unchanged; for the analyses reported here all selected positions were masked.

Multi-sampled PPPL extended the sampled variant by generating multiple sampled-mask realizations of the same sequence in a single batch (8 realizations in the reported runs). Selected positions were allocated row-wise so that every valid sequence position appeared as a masked position in at least one realization, guaranteeing complete coverage at finite batch sizes. Per-position losses were averaged across the realizations that covered each position before exponentiating the mean.

Direct PPPL replaced the masked inference loop with a trained regression head *f_φ_* that predicts per-position masked losses from the ESM-2 final-layer hidden states in a single forward pass,

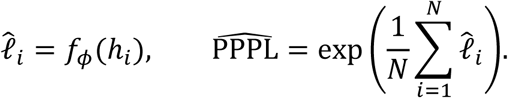

This reduced PPPL inference complexity from O(N) forward passes to a single forward pass per sequence.

### Reference k-mer Encoding for Positional Analyses

To distinguish residues found in a curated reference proteome from candidate-novel residues, we annotated each input sequence by its k-mer coverage against a precomputed reference k-mer set of length k =9. The reference set was built once from the comprehensive database used in our mass-spectrometry pipeline, which combines the UniProt human reference proteome (UP000005640, 2026_05_2 release) with Gencode v42 protein-coding non-redundant sequences, supplementary catalogs of small ORFs (smORFs, n = 2,043) and non-canonical ORFs (nuORF v1), 602 common laboratory contaminants, and JPT iRT calibration peptides. Every 9-mer substring of every sequence in this database was enumerated and stored as a Python set, yielding the reference k-mer set used downstream.

For each input position we determined whether it fell inside at least one k-mer present in the reference set (“reference-covered”) and whether it fell inside at least one k-mer absent from the reference set (“unique”). Positions were then encoded into a per-residue annotation string with three possible states: positions covered only by unique (non-reference) k-mers retained the original uppercase amino-acid letter; positions covered by both unique and reference k-mers were rewritten in lowercase; and positions covered by no unique k-mer were replaced by X. Per-sequence masked counts and percentages were stored alongside the encoded sequence for downstream filtering.

### Token-Regression Head: Training Data

The direct PPPL regression head was trained on a token-loss dataset constructed by labeling sampled reference proteins with their standard masked per-position losses. Starting from a human lung cryptic proteome dataset, sequences longer than 1024 residues were filtered out, leaving 17,881,596 candidates. A uniform random sample of 10,000 sequences was drawn at seed 42, masked PPPL was computed for each at the 650M backbone with batch size 8, and per-position token losses and token identifiers were saved as a Hugging Face DatasetDict. The dataset was split at seed 42 into 9000 / 500 / 500 train / validation / test partitions.

### Token-Regression Head: Architecture

The regression head attached an ensemble of feed-forward heads to the frozen final hidden states of the ESM-2 backbone. Each individual head consisted of depth repeated blocks of (Linear → LayerNorm → GELU → Dropout) followed by a scalar output projection, and the ensemble output was the unweighted mean of the per-head scalar predictions. The reported 650M run used 8 heads of depth 3, hidden dropout 0.1, and shared the backbone hidden size. The ESM-2 backbone was held frozen throughout training; only the regression-head parameters were updated.

### Token-Regression Head: Training Procedure

The head was trained with the Hugging Face Trainer using a per-token mean-squared-error objective. The loss at each position was the squared difference between the predicted scalar and the labeled masked loss, restricted to real amino-acid positions (special tokens, padding, and any other positions without a labeled loss were excluded). Per-sequence losses were averaged within sequence, and the batch loss was averaged across sequences containing at least one valid target position. Training used a learning rate of 1×10⁻⁴, a per-device batch size of 16 (evaluation batch size 64), a maximum sequence length of 1024, and ran for 50 epochs. Evaluation and checkpoint logging were performed every 100 steps, and the best checkpoint was selected on minimum validation pppl_rmse.

Validation PPPL metrics were reconstructed at the sequence level. For each validation sequence the predicted and true per-token losses were averaged over valid positions and exponentiated to give predicted and true PPPL, and the mean absolute error, root-mean-squared error, and coefficient of determination were computed across sequences. The 650M-backbone training run selected its best checkpoint at global step 24,200, with validation pppl_rmse = 0.6541, pppl_r2 = 0.9925, and pppl_mae = 0.3294. This checkpoint was used as *f_φ_* for all subsequent cohort-scale analyses.

### Sequence Perturbation Benchmark

To characterize PPPL behavior across model sizes and perturbation severity we evaluated all five PPPL protocols on the same Cytochrome P450 3A4 (CP3A4_HUMAN) reference protein and a battery of derived controls: the unmodified original sequence; a uniformly shuffled sequence with the start methionine preserved; a uniformly random amino-acid sequence of identical length; the first 250-residue truncation; a block-shuffled version in which the sequence beyond the start methionine was partitioned into random-length blocks of 25–50 residues and the blocks reordered; three point-mutated variants at 15%, 25%, and 50% per-position substitution probability; and a same-length translation of a pseudogene locus serving as a biologically meaningful intermediate control.

### Cohort-Scale Direct Scoring

For population-level analyses we computed direct PPPL on two protein cohorts: (1) a high-confidence set of cryptic protein translation products predicted by our internal RyboCypher pipeline, n = 3,192 representative sequences after MMseqs2 clustering at 99% sequence identity; and (2) a human reference set drawn from peptide-supported UniProt human proteins, n = 1,017 representative sequences after the same 99%-identity clustering.

Both cohorts were processed identically: each input FASTA was clustered at 99% identity (MMseqs2 easy-cluster, --cluster-mode 2 --cov-mode 1), the representative sequences were scored at the 650M backbone with ppl_type=’direct’ and batch_size = 8, and the reference k-mer DB (k = 9, diff split mode) was applied to produce per-sequence masked counts, masked percentages, masked-position geometric-mean PPPL, and unmasked-position geometric-mean PPPL.

### Statistical Analyses

Cohort-level distributional comparisons were performed on the raw direct PPPL values and on the masked-position and unmasked-position geometric-mean PPPL values, separately for four filter regimes: unfiltered; length > 100 residues; 90%-identity cluster representatives only; and the conjunction of the latter two. Group differences were summarized using the mean, median, and standard deviation per cohort, and tested using the two-sided Mann–Whitney U test on the underlying values. Effect sizes were reported as Cliff’s delta computed from the Mann–Whitney U statistic. Positional analyses for individual selected proteins compared the geometric mean of predicted PPPL within the masked and unmasked residue groups defined by the reference k-mer encoding.

### Cryptic ORF ESMFold structure prediction

Full-length protein sequences for three cryptic ORF candidates identified by mass spectrometry were folded using ESMFold2^61^ (Biohub, 2026). ESMFold2 predicts three-dimensional structure from primary sequence alone using a 6-billion parameter protein language model with a diffusion-based structure module, requiring no multiple-sequence alignment or homology templates. Per-residue confidence scores (pLDDT, 0 to 100) were extracted from the B-factor column of each output PDB file and used to assess local model confidence. The position of each MS-detected peptide within its parent ORF was determined by exact amino acid string matching against the full protein sequence. Structures were visualized and figures rendered using py3Dmol (v2.5.5) in Python 3.12.

## Antibody development

### pMHC Targets

*In vitro* targets were produced using easYmers® (Immunodex), according to the manufacturer’s instructions. Briefly, peptide targets were synthesized alongside controls and exchanged into receptive MHC monomers for HLA-A*02:01.

### Antibody Library (scFv) Screening and Affinity Maturation and Cell Binding

Antibodies were generated by successive selection of large human single-chain variable fragment (scFv) libraries by phage display. Affinity for pMHC target were determined by SPR and binding to control cell lines. Top candidates were affinity matured and further tested for HLA haplotype restriction, using HEK293T cells expressing the target pMHC formatted for either HLA A*2:01, A*2:05 and A*2:06 with the target peptide or controls. Binding was determined by staining cells with full length antibody, with detection using a labeled secondary antibody. Shift in mean fluorescence intensity (MFI) was used to determine affinity.

### ADC Conjugations and In vitro Cell Killing

Antibody drug conjugates (ADC) were produced using maleimide directed chemistry to install Exatecan and Eribulin toxic payloads connected to a full-length antibody using a cleavable linker. Cell viability was performed using CellTiter-Glo® 2.0 Cell Viability Assay (Promega), according to the manufacturer’s instructions. Briefly, HCC78 cells were plated in white multi-well plates in appropriate treatment conditions, cultured and equal volumes of viability reagent was added to each well with luminescence acquired in a Varioskan LUX Multimode Microplate Reader (Thermo Scientific).

