## Supplemental Information for "Discovery and Targeting of a Cryptic Human Proteome"

### Supplementary Fig. 1

a

| Cancer Type | Label | Unique RNA Loci | Unique Isoforms | Samples | % Total Loci |
|---|---|---|---|---|---|
| CRC Tumors | CRC | 1,024,598 | 1,181,424 | 3 | 15.5% |
| NSCLC Tumors | NSCLC | 3,598,701 | 4,817,182 | 8 | 54.5% |
| NSCLC Healthy | NSCLC HEALTHY | 2,287,241 | 2,635,974 | 3 | 34.6% |
| NSCLC Cancer Cell Line | CCL | 105,891 | 136,959 | 1 | 1.6% |
| **Total** | — | **6,605,037** | **8,308,621** | **15** | — |

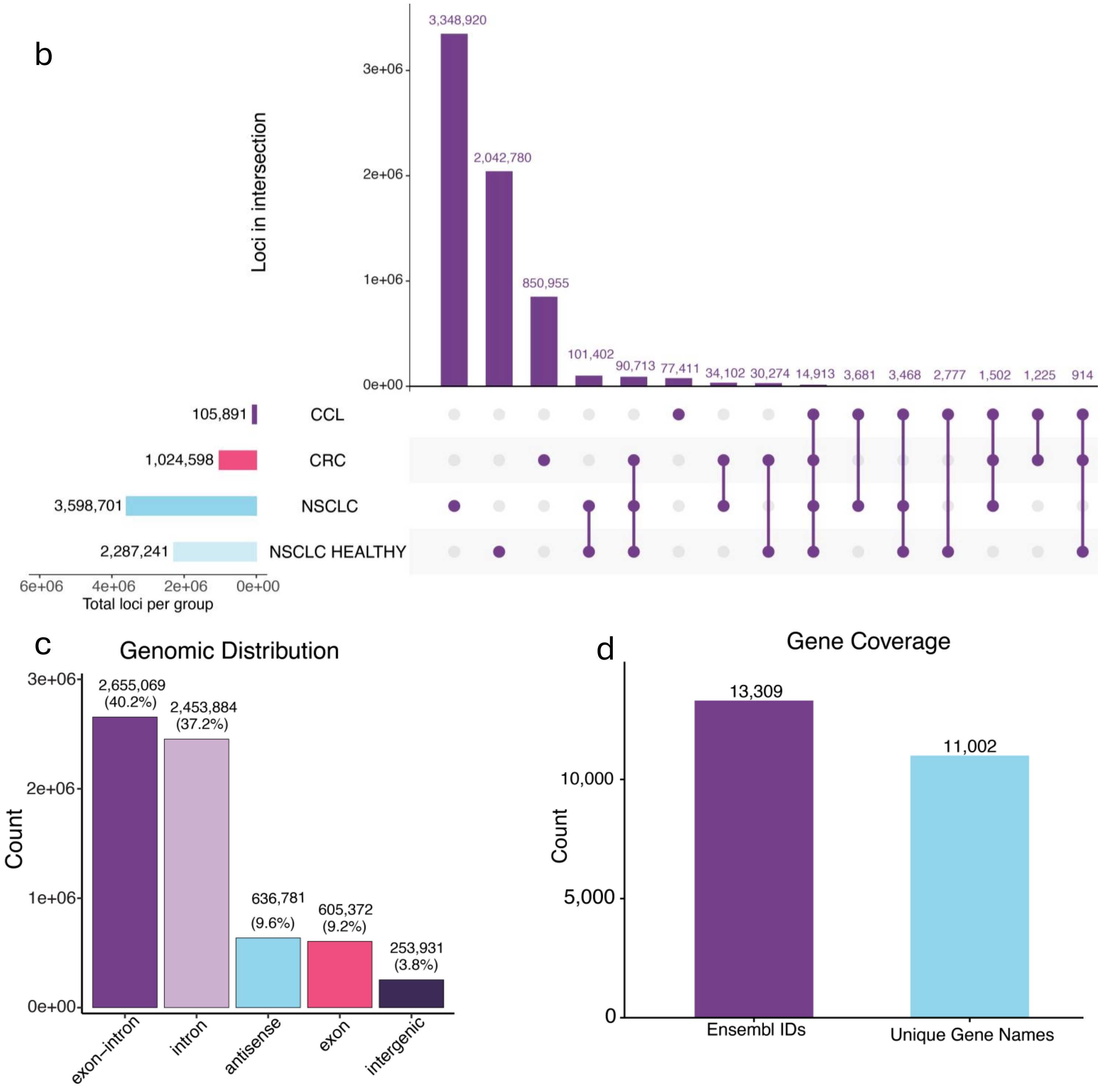

**Supplementary Fig. 1 | RyboCypher dark RNA characterization.**

**(a)** Summary table of unique RNA loci and isoforms per sample group: CRC Tumors (1,024,598 loci; 1,181,424 isoforms; n = 3; 15.5%), NSCLC Tumors (3,598,701; 4,817,182; n = 8; 54.5%), NSCLC Healthy (2,287,241; 2,635,974; n = 3; 34.6%), NSCLC Cancer Cell Line (105,891; 136,959; n = 1; 1.6%); total 6,605,037 loci / 8,308,621 isoforms across 15 samples. **(b)** Upset plot of loci intersections across sample groups. Largest single-group intersection: NSCLC tumors (3,348,920), followed by shared NSCLC/NSCLC-Healthy loci (2,042,780) and CRC-exclusive loci (850,955). **(c)** Genomic distribution of retained loci: exon-intron junctions (2,655,069; 40.2%), intronic (2,453,884; 37.2%), antisense (636,781; 9.6%), exonic (605,372; 9.2%), intergenic (253,931; 3.8%). **(d)** Gene coverage: 13,309 Ensembl IDs corresponding to 11,002 unique gene names.

**Supplementary Table 1 | CPTAC Cellular and Membrane Proteomics Datasets**[29,32-43]

| CPTAC Accession | Study Name | Cancer Type | Patients |
|---|---|---|---|
| PDC000153 [29] | CPTAC LUAD Discovery Study - Proteome | Lung Adenocarcinoma | 115 |
| PDC000489 [40] | CPTAC LUAD Confirmatory Study - Proteome | Lung Adenocarcinoma | 131 |
| PDC000118 [41] | Prospective Ovarian PNNL Proteome Qeplus | Ovarian Serous Cystadenocarcinoma | 95 |
| PDC000127 [32] | CPTAC CCRCC Discovery Study - Proteome | Clear Cell Renal Cell Carcinoma;Non-Clear Cell Renal Cell Carcinoma | 124 |
| PDC000116 [33] | Prospective Colon PNNL Proteome Qeplus | Colon Adenocarcinoma | 102 |
| PDC000125 [34] | CPTAC UCEC Discovery Study - Proteome | Uterine Corpus Endometrial Carcinoma | 123 |
| PDC000234 [40] | CPTAC LSCC Discovery Study - Proteome | Lung Squamous Cell Carcinoma | 115 |
| PDC000614 [35,42] | CPTAC STAD Study - Proteome | Early Onset Gastric Cancer, Stomach Adenocarcinoma | 193 |
| PDC000221 [36] | CPTAC HNSCC Discovery Study - Proteome | Head and Neck Squamous Cell Carcinoma | 124 |
| PDC000270 [37] | CPTAC PDA Discovery Study - Proteome | Pancreatic Ductal Adenocarcinoma | 166 |
| PDC000606 [38] | SIMM Gallbladder Cancer - Proteome | Epithelial Neoplasms | 195 |
| PDC000445 [34] | CPTAC UCEC Confirmatory Study - Glycoproteome | Uterine Corpus Endometrial Carcinoma | 159 |
| PDC000272 [37] | CPTAC PDA Discovery Study - Intact Glycoproteome | Pancreatic Ductal Adenocarcinoma | 166 |
| PDC000250 [39] | Prospective Ovarian JHU Intact Glycoproteome | Ovarian Serous Cystadenocarcinoma | 97 |
| PDC000471 [43] | CPTAC CCRCC Discovery Study - Intact Glycoproteome | Clear Cell Renal Cell Carcinoma; Non-Clear Cell Renal Cell Carcinoma | 124 |

**Supplementary Table 2 | Publicly Available pMHC Datasets**[44-54]

| Study | Cancer Type | Patients |
|---|---|---|
| Nelde et al [52] | Acute lymphocytic leukemia | 47 |
| Huwiler et al [53] | Acute Myeloid leukemia | 8 |
| Apavaloaei et al [45] | Lung Adenocarcinoma | 12 |
| Kraemer et al [48] | Lung Adenocarcinoma | 8 |
| Kina et al [46] | Breast cancer | 24 |
| Bassani-Sternberg et al [44] | Melanoma | 19 |
| Ely et al [51] | Pancreatic Ductal Adenocarcinoma (organoids) | 11 |
| Ehx et al [47] | Acute myleoid Leukemia | 14 |
| Newey et al [50] | Colon (Organoids) | 6 |
| Raja et al [54] | Ovarian | 5 |

### Supplementary Fig. 2

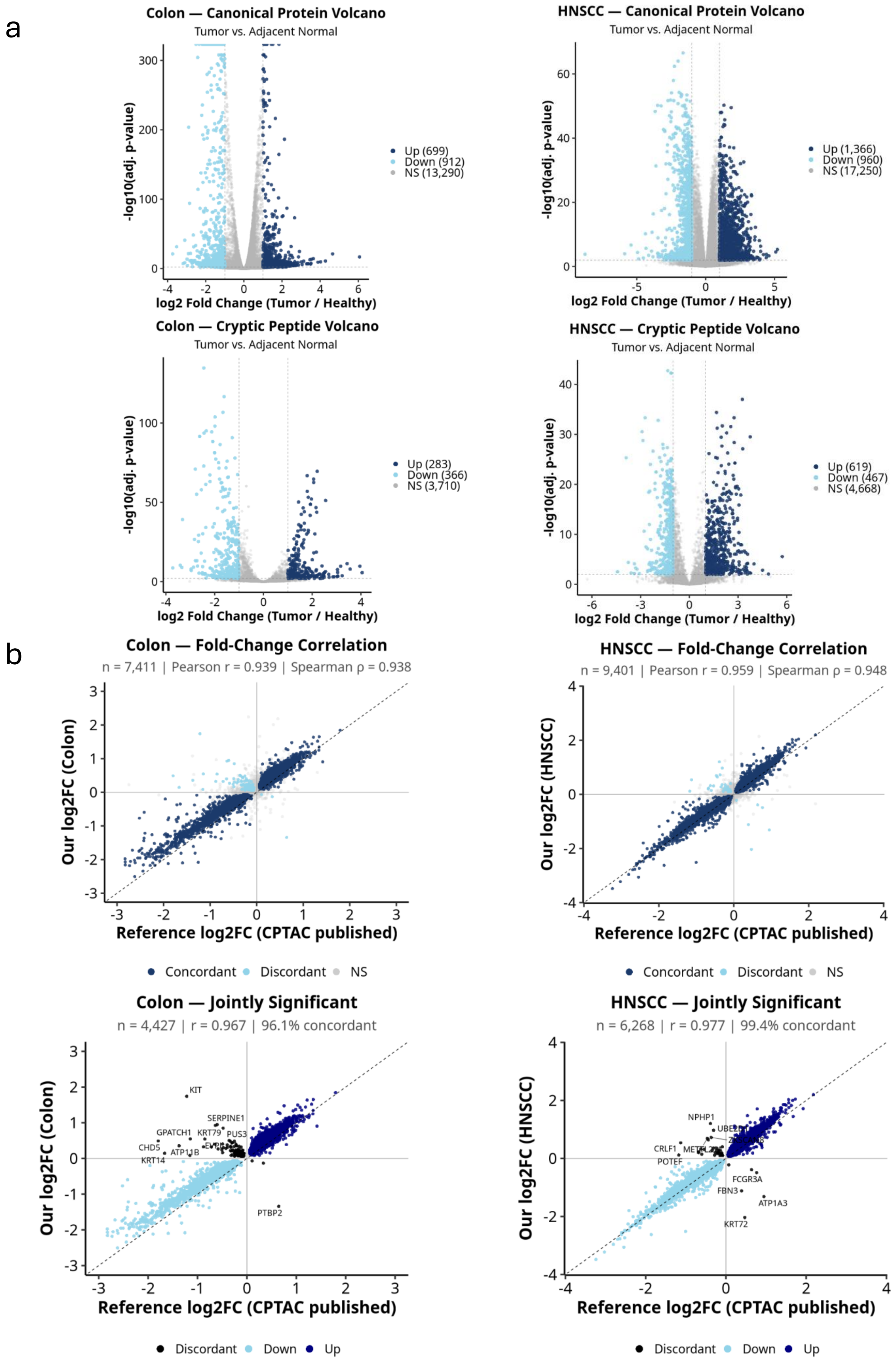

**Supplementary Fig. 2 | Discovery and initial characterization of a cryptic human proteome.**

**(a)** DarkRNA-encoded cryptic peptidome landscape across CPTAC cellular proteomic cohorts. Paired volcano plots of canonical protein (top row) and cryptic peptide (bottom row) differential abundance for Colon and HNSCC (Tumor vs. Adjacent Normal; adj. $p < 0.01$, $|log2FC| >= 1$). Colon canonical: Up (699), Down (912), NS (13,290); HNSCC canonical: Up (1,366), Down (960), NS (17,250). Colon cryptic: Up (283), Down (366), NS (3,710); HNSCC cryptic: Up (619), Down (467), NS (4,668). **(b)** Per-cohort cross-validation of canonical protein log2FC vs. published CPTAC reference for Colon and HNSCC. Top row: full fold-change correlations (Colon: $n = 7,411$, Pearson $r = 0.939$, Spearman $r = 0.938$; HNSCC: $n = 9,401$, Pearson $r = 0.959$, Spearman $r = 0.948$); bottom row: jointly significant subset with discordant outliers labeled (Colon: $n = 4,427$, $r = 0.967$, 96.1% concordant; HNSCC: $n = 6,268$, $r = 0.977$, 99.4% concordant).

### Supplementary Fig. 3

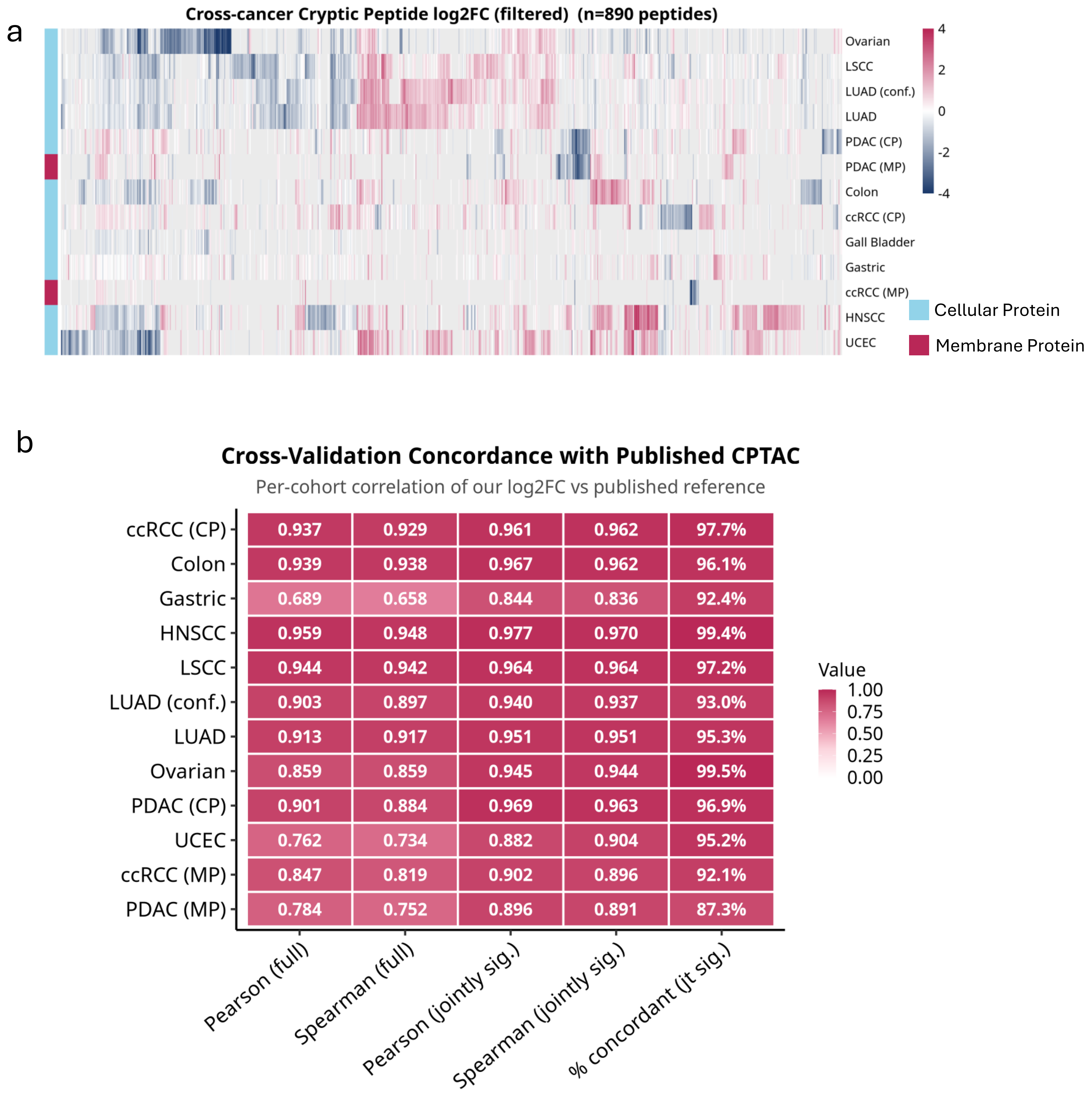


**Supplementary Fig. 3 | Initial characterization of a cryptic human proteome.**

**(a)** Cross-cancer cryptic peptide log2FC heatmap (filtered; n = 890 peptides; adj. p < 0.01, |log2FC| >= 1). Cohorts: Ovarian, LSCC, LUAD (conf.), LUAD, PDAC (CP), PDAC (MP), Colon, ccRCC (CP), Gallbladder, Gastric, ccRCC (MP), HNSCC, UCEC. Colored by assay: Cellular Protein (blue), Membrane Protein (pink). Rows hierarchically clustered; columns ordered by cohort. **(b)** Cross-validation concordance with published CPTAC. Per-cohort correlation of log2FC vs. published reference across 12 cohorts: ccRCC CP (Pearson 0.937, Spearman 0.929, jointly sig. Pearson 0.961, Spearman 0.962, 97.7% concordant), Colon (0.939, 0.938, 0.967, 0.962, 96.1%), Gastric (0.689, 0.658, 0.844, 0.836, 92.4%), HNSCC (0.959, 0.948, 0.977, 0.970, 99.4%), LSCC (0.944, 0.942, 0.964, 0.964, 97.2%), LUAD conf. (0.903, 0.897, 0.940, 0.937, 93.0%), LUAD (0.913, 0.917, 0.951, 0.951, 95.3%), Ovarian (0.859, 0.859, 0.945, 0.944, 99.5%), PDAC CP (0.901, 0.884, 0.969, 0.963, 96.9%), UCEC (0.762, 0.734, 0.882, 0.904, 95.2%), ccRCC MP (0.847, 0.819, 0.902, 0.896, 92.1%), PDAC MP (0.784, 0.752, 0.896, 0.891, 87.3%).

#### Supplementary Fig. 4

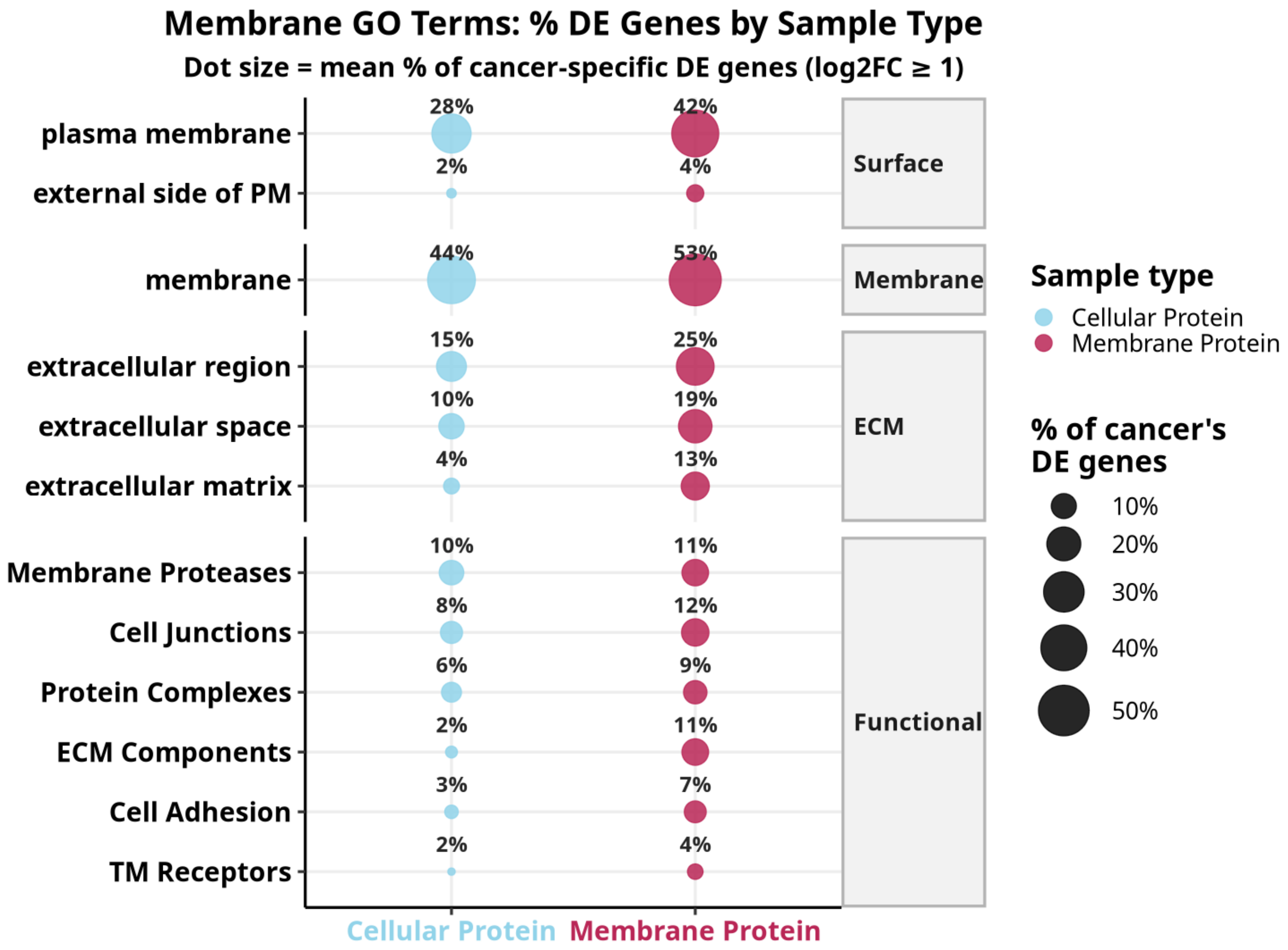


**Supplementary Fig. 4 | Selection of actionable membrane targets from the cryptic cancer proteome.**
Membrane GO Terms: mean percentage of cancer-associated DE genes (log2FC >= 1) by sample type. Dot size encodes mean percentage across cancers; Cellular Protein (light blue), Membrane Protein (pink). Surface terms: plasma membrane (28% Cellular, 42% Membrane), external side of PM (2%, 4%). Membrane: 44% Cellular, 53% Membrane. ECM terms: extracellular region (15%, 25%), extracellular space (10%, 19%), extracellular matrix (4%, 13%). Functional terms: Membrane Proteases (10%, 11%), Cell Junctions (8%, 12%), Protein Complexes (6%, 9%), ECM Components (2%, 11%), Cell Adhesion (3%, 7%), TM Receptors (2%, 4%). Membrane Protein consistently enriches surface-accessible and ECM-associated categories at higher fractions than Cellular Protein, supporting the use of membrane proteomics for identifying cryptic targets with extracellular accessibility.

### Supplementary Fig. 5

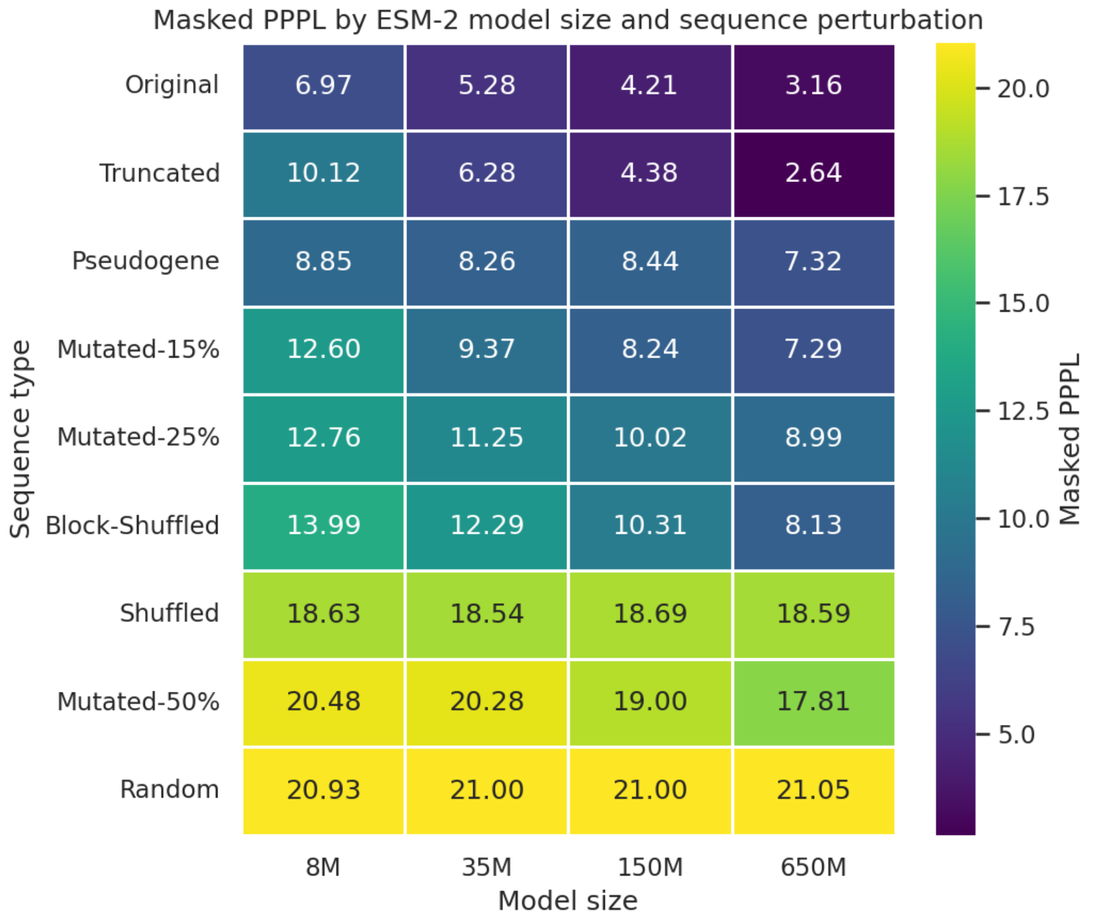


**Supplementary Fig. 5 | Pseudo-perplexity tracks perturbation severity across ESM-2 model sizes.** Standard masked pseudo-perplexity (PPPL) for the canonical reference protein CP3A4_HUMAN and a graded panel of synthetic perturbations (an N-terminal truncation, a block shuffle, point mutations at 15%, 25%, and 50%, a fully shuffled sequence, a uniformly random sequence, and a same-length pseudogene-derived translation) evaluated at four ESM-2 backbones (8M, 35M, 150M, and 650M parameters). Rows are sorted by mean PPPL; lower values indicate sequences closer to natural protein space. Mean PPPL increases monotonically with perturbation severity, from 4.9 for the unperturbed sequence to 21.0 for the uniformly random control, with the pseudogene-derived translation at an intermediate value of 8.2, and the ordering is stable across model sizes.

### Supplementary Fig. 6

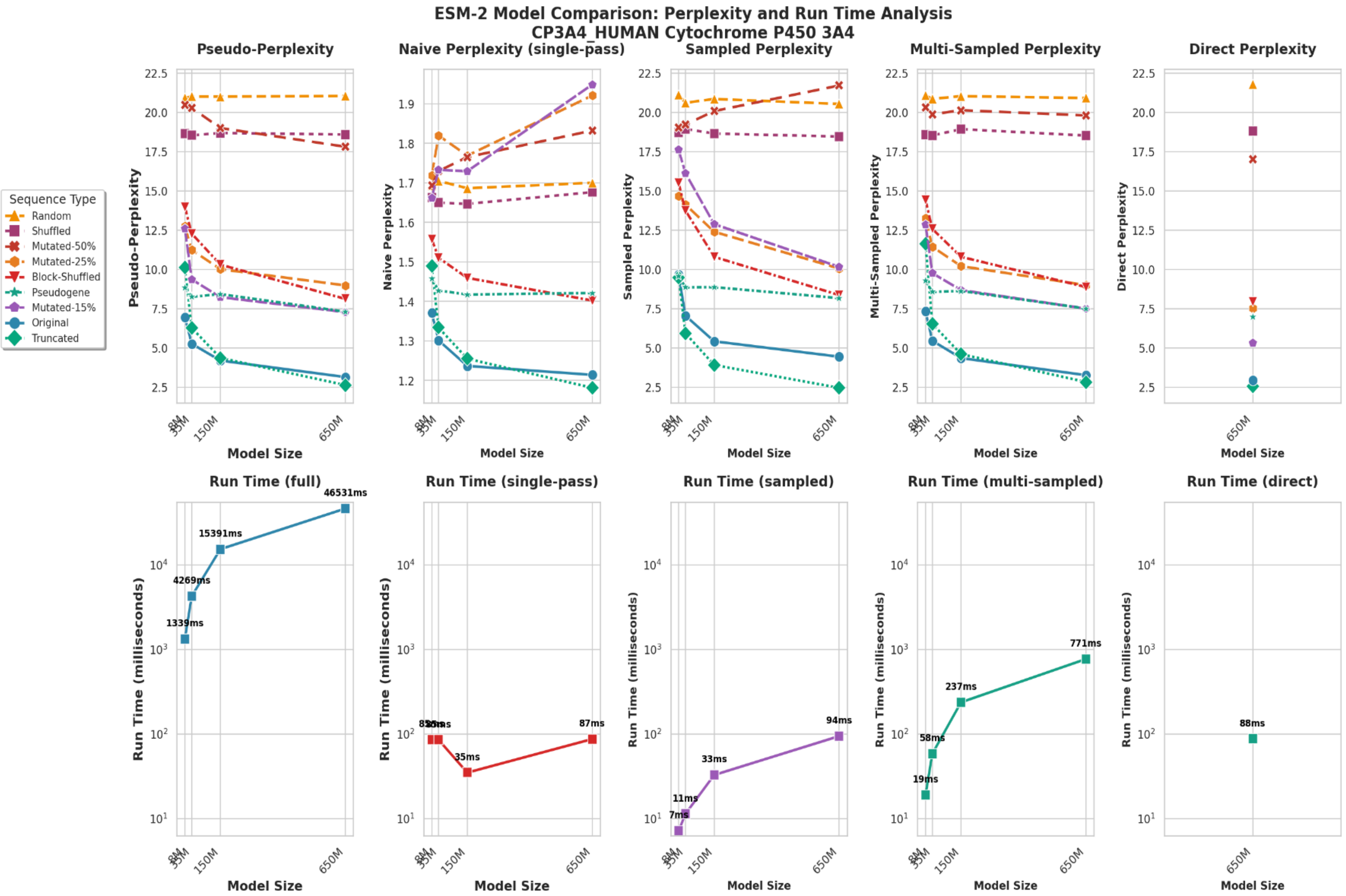


**Supplementary Fig. 6 | Speed–fidelity comparison of five PPPL protocols.**
Five PPPL protocols (standard masked, naive, sampled, multi-sampled, and direct) were evaluated on the nine-class sequence-perturbation benchmark across the four ESM-2 model sizes. Top row, PPPL versus model size (one line per sequence class); bottom row, mean inference run time (logarithmic scale). Standard masked PPPL separates the sequence classes cleanly with an ordering stable across model sizes; naive PPPL collapses to a narrow, non-discriminative band; sampled and multi-sampled PPPL preserve the masked ordering with progressively lower variance; and direct PPPL reproduces the masked ordering at single-pass cost.

### Supplementary Fig. 7

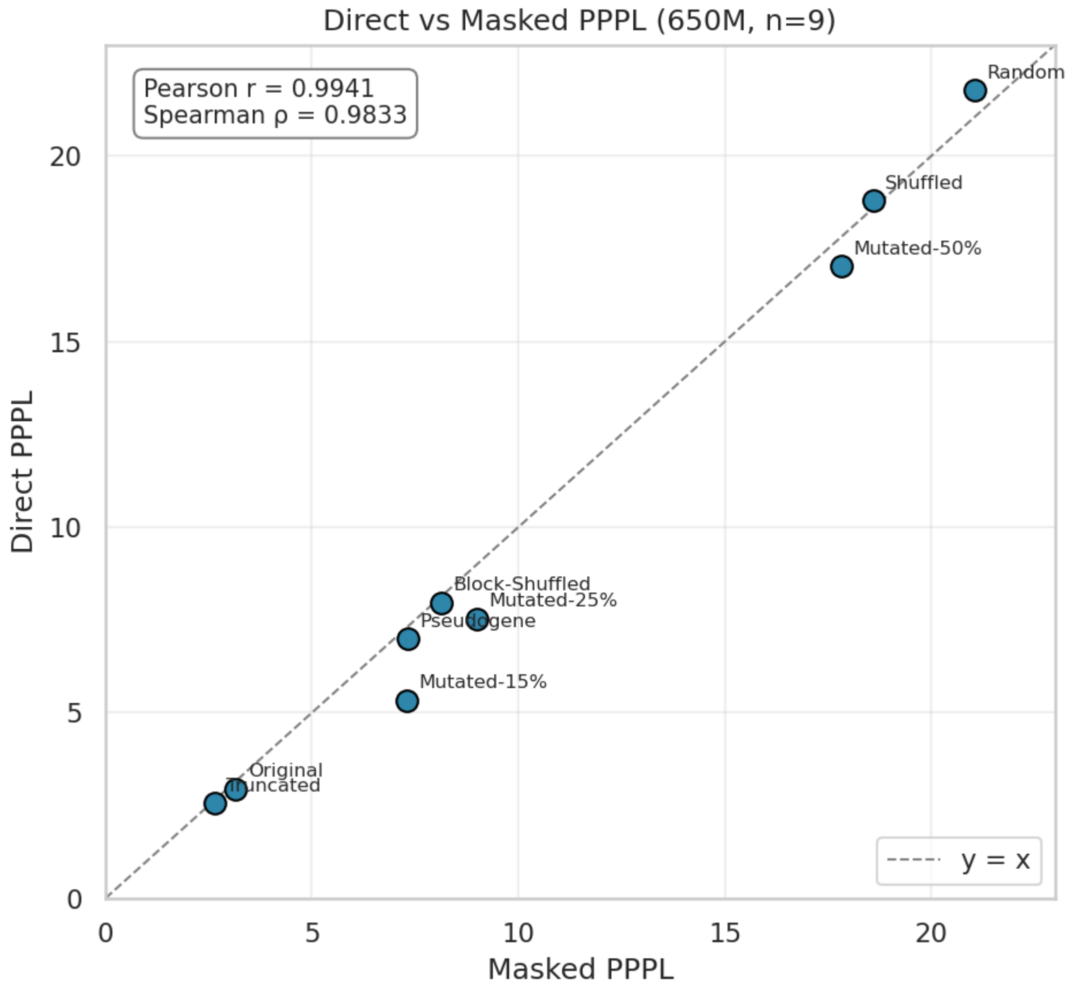


**Supplementary Fig. 7 | Direct PPPL reproduces standard masked PPPL.**
Direct versus standard masked PPPL across the nine sequence classes at the 650M ESM-2 backbone; the dashed line is the y = x identity. Direct and masked PPPL agree closely across an approximately seven-fold range of PPPL (Pearson r = 0.994, Spearman ρ = 0.983), with the largest discrepancies confined to the mid-perturbation regime.

### Supplementary Fig. 8

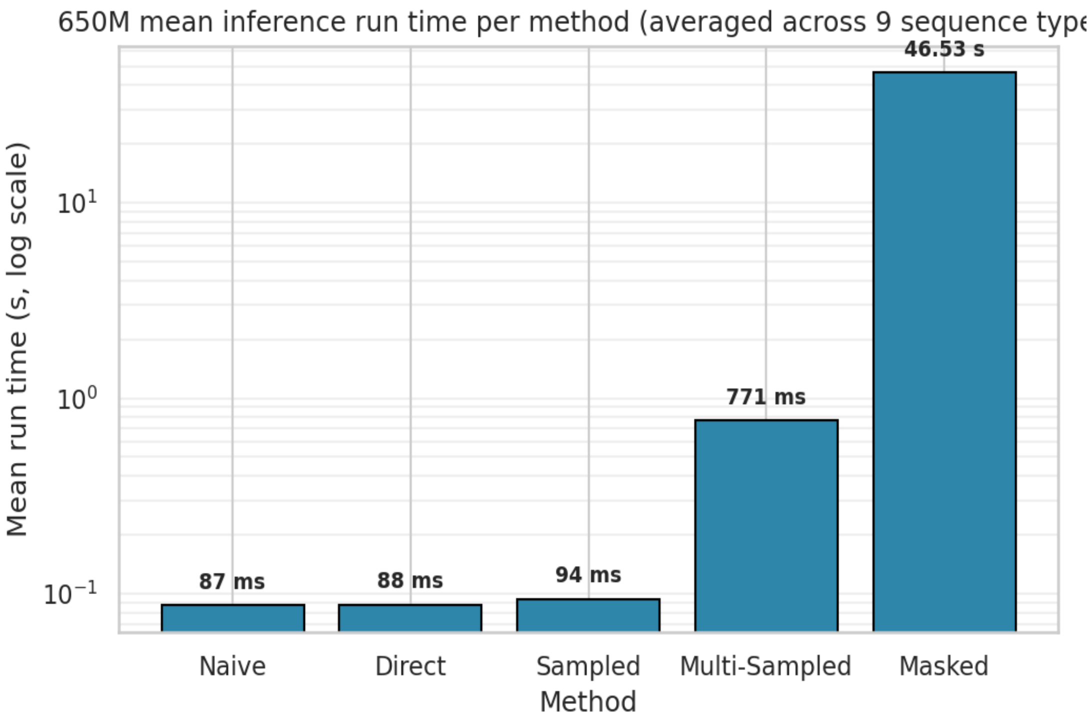


**Supplementary Fig. 8 | Direct PPPL is orders of magnitude faster than masked PPPL.**
Mean inference run time per PPPL protocol at the 650M ESM-2 backbone, averaged across the nine sequence classes (logarithmic y-axis). Direct PPPL averages 88 ms per sequence versus 46.5 s for standard masked PPPL (an approximately 530-fold reduction) while preserving the fidelity shown in Supplementary Fig. 7.

#### Supplementary Fig. 9

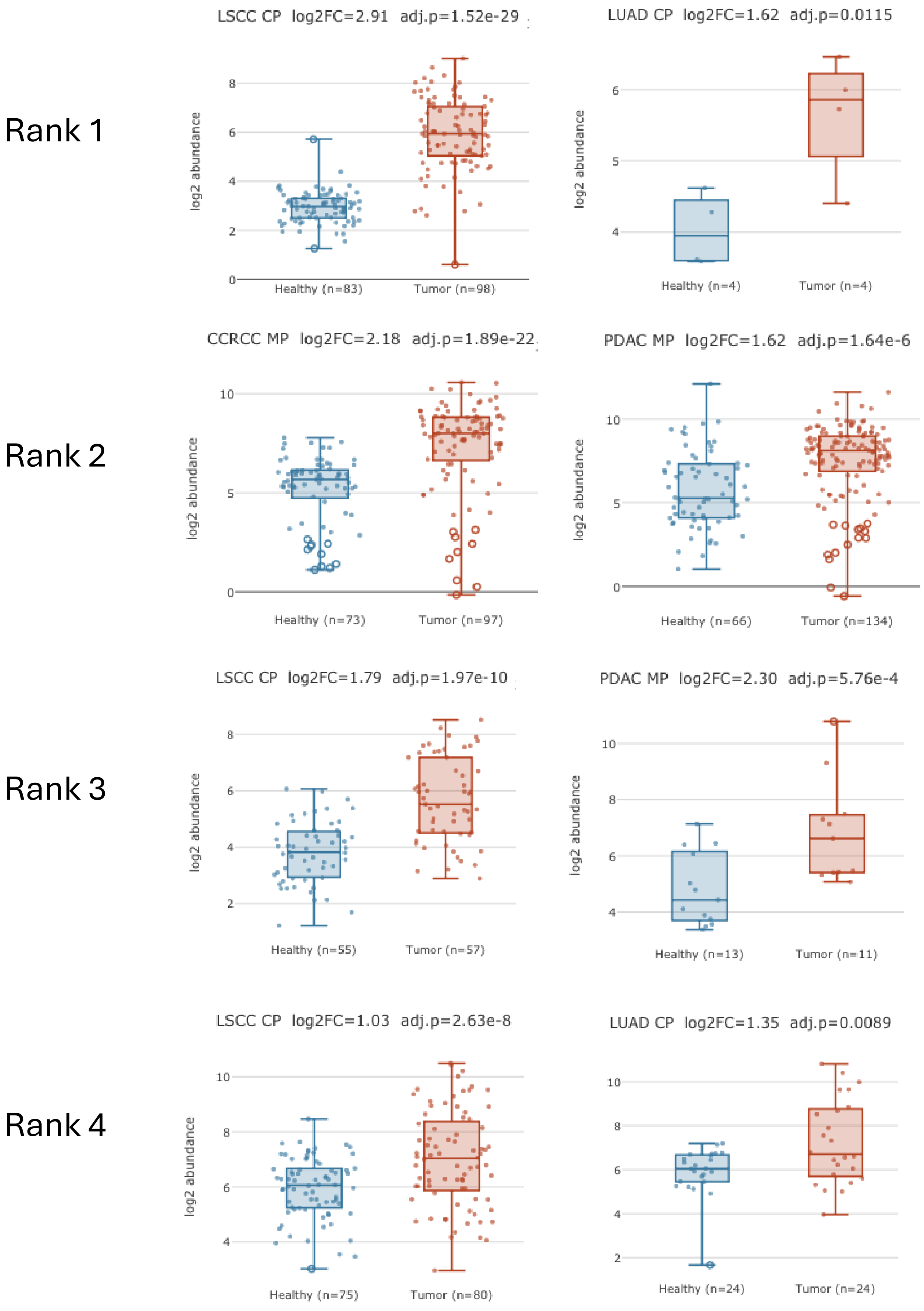

**Supplementary Fig. 9 | Differential expression of peptides mapping to top ranked cryptic membrane proteins.**
Tumor vs. healthy log2 abundance boxplots for the top four ranked membrane target cryptic ORFs from Fig. 4a, shown across their most differentially expressed CPTAC cohorts. Rank 1 (RD-8886583): Lung Squamous Cell Carcinoma (LSCC) cellular proteomics log2FC = 2.91, adj. p = 1.52e-29 (Healthy n = 83, Tumor n = 98); Lung Adenocarcinoma (LUAD) cellular proteomics log2FC = 1.62, adj. p = 0.0115 (Healthy n = 4, Tumor n = 4); Clear Cell Renal Cell Carcinoma (ccRCC) membrane proteomics log2FC = 2.18, adj. p = 1.89e-22; Pancreatic Ductal Adenocarcinoma (PDAC) membrane proteomics log2FC = 1.62, adj. p = 1.64e-6. Rank 2 (RD-5336591): LSCC cellular proteomics (Healthy n = 73, Tumor n = 97); PDAC membrane proteomics (Healthy n = 66, Tumor n = 134). Rank 3 (RD-0376778): LSCC cellular proteomics log2FC = 1.79, adj. p = 1.97e-10 (Healthy n = 55, Tumor n = 57); PDAC membrane proteomics log2FC = 2.30, adj. p = 5.76e-4 (Healthy n = 13, Tumor n = 11). Rank 4 (RD-8653598): LSCC cellular proteomics log2FC = 1.03, adj. p = 2.63e-8 (Healthy n = 75, Tumor n = 80); LUAD cellular proteomics log2FC = 1.35, adj. p = 0.0089 (Healthy n = 24, Tumor n = 24). Blue = healthy, red = tumor.

### Supplementary Fig. 10

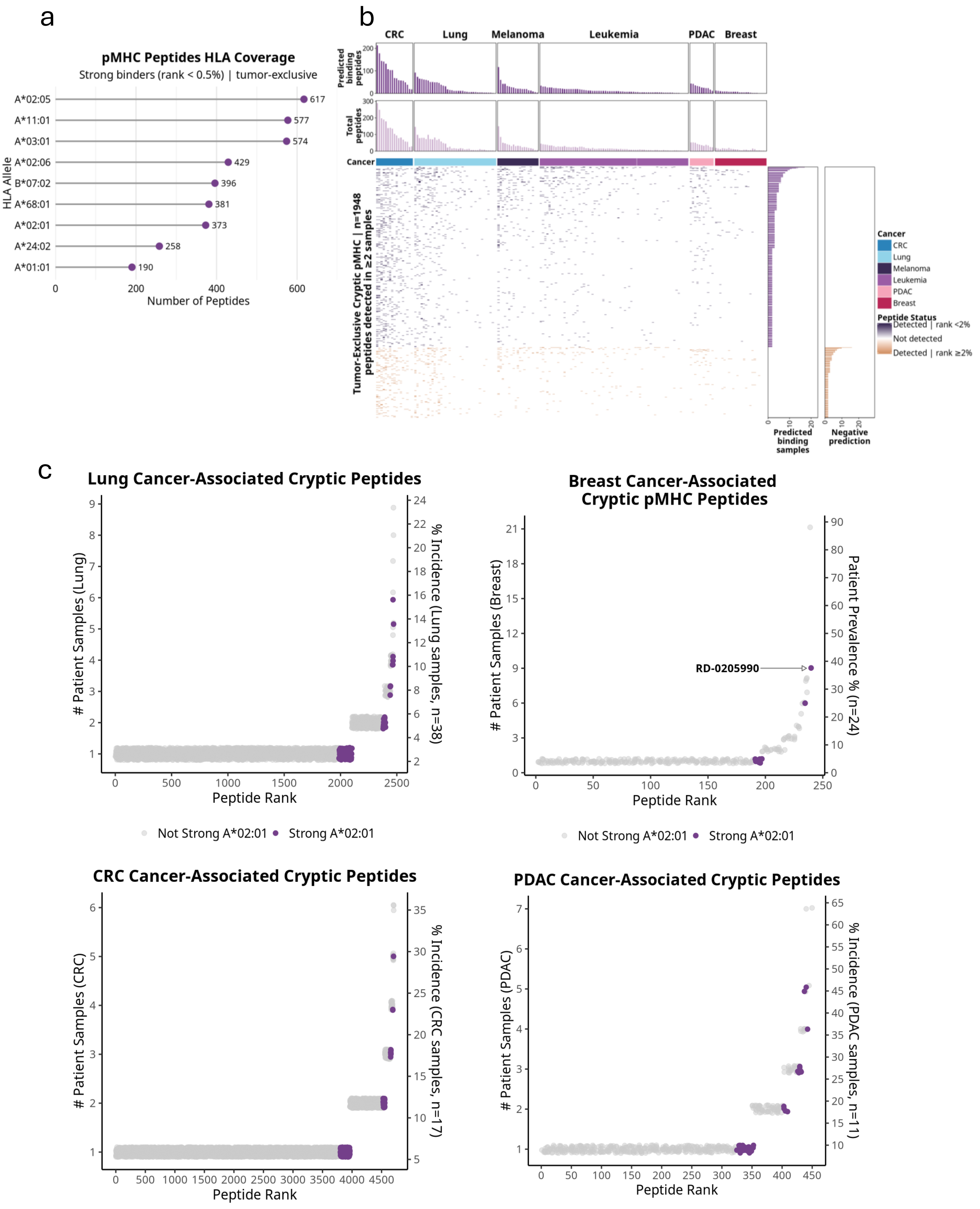

**Supplementary Fig. 10 | Selection of actionable targets from the cryptic cancer immunopeptidome - extended data.**

**(a)** pMHC Peptides HLA Coverage: tumor-exclusive cryptic strong binders (MHCflurry rank <0.5%) per HLA allele. A*02:05 (617), A*11:01 (577), A*03:01 (574), A*02:06 (429), B*07:02 (396), A*68:01 (381), A*02:01 (373), A*24:02 (258), A*01:01 (190). **(b)** Per-sample detection matrix of tumor-exclusive cryptic pMHC peptides detected in >=2 samples (n = 1,948 peptides) across CRC, Lung, Melanoma, Leukemia, PDAC, and Breast. Purple = detected, rank <2%; orange = detected, rank >=2%; grey = not detected. **(c)** Ranked patient prevalence scatter plots for tumor-exclusive HLA-A*02:01 strong binder cryptic pMHC peptides in Lung (n = 38), Breast (n = 24), CRC (n = 17), and PDAC (n = 11). Purple = strong A*02:01 binder; grey = not strong. RD-0205990 annotated.

#### Supplementary Fig. 11

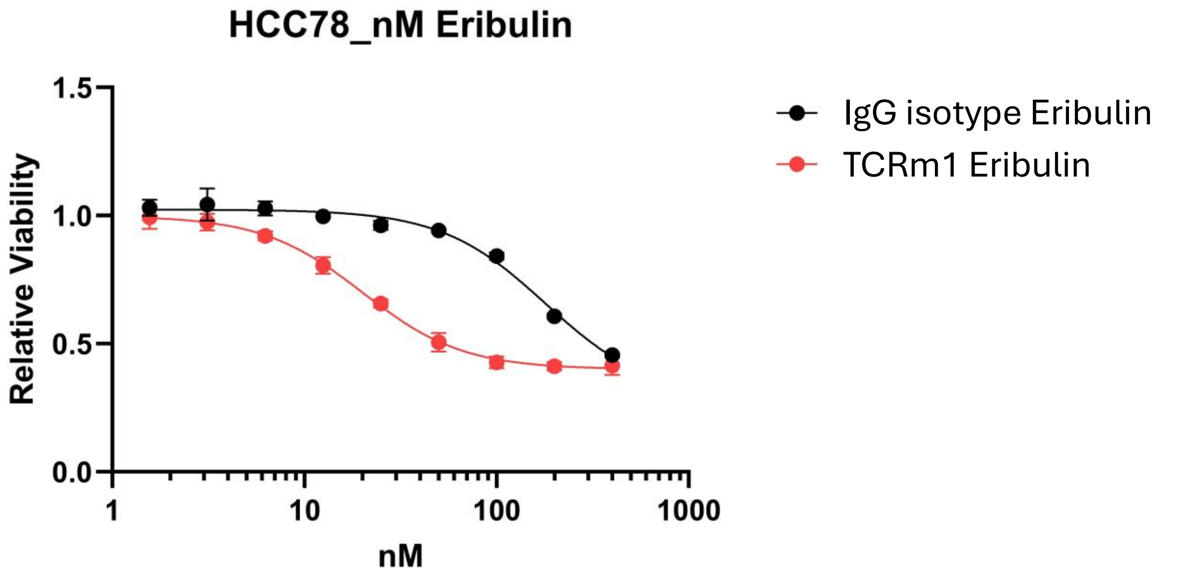


| | IC50 | R squared |
|---|---|---|
| TCRm1 Eribulin | 20.06 | 0.9881 |

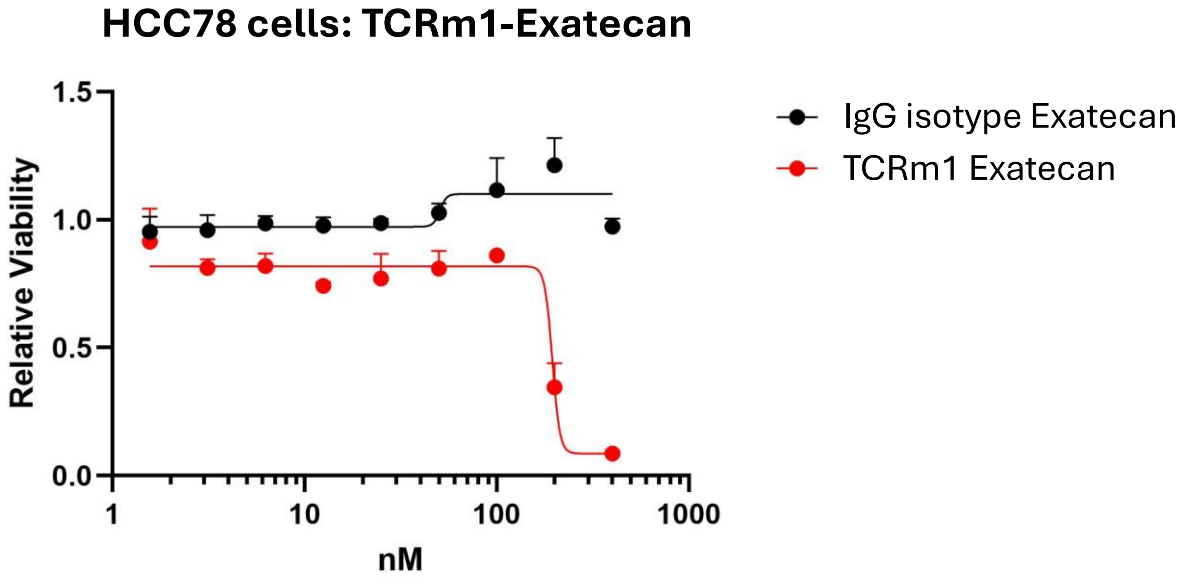


| | IC50 | R squared |
|---|---|---|
| TCRm1 Exatecan | 194.7 | 0.9259 |

**Supplementary Fig. 11 | Demonstration of *in vitro* cancer cell killing by TCRm1 formatted as ADCs.**
**(Top)** TCRm1 was reduced with TCEP, and Eribulin was conjugated via maleimide to free sulfhydryl residues (cysteines) through a lysosome-cleavable linker, to arrive at an average drug to antibody ratio (DAR) of ~6 (red). The same reaction conditions were used for linker/payload (L/P) derivatization of an anti-beta galactosidase human IgG1 antibody (IgG isotype control-Eribulin (black)). HCC78 cells were plated (3e3 cells/well) in 96 well plates and treated with a serial dilution of antibody (400, 200, 100, 50, 25, 12.5, 6.25, 3.12, and 1.56nM). Plates were incubated for 84h before cell viability determination was performed with CelTiter-Glo 2.0. Exposure of cells to 50nM TCRm1-Eribulin achieved a relative viability score of 0.5 in this assay (calculated $IC_{50}$ ~20nM), while the IgG isotype control-Eribulin conjugate approached this value only at the highest concentrations tested (200 and 400nM). **(Bottom)** TCRm1 was conjugated to a maleimide-functionalized, lysosome-cleavable linker to Exatecan under the same reaction conditions used for derivatization with Eribulin, with slightly lower DAR achieved (~4.5 (red)). The Exatecan L/P was also conjugated to the same anti-beta galactosidase human IgG1 antibody (IgG isotype control-Exatecan (black)). HCC78 cells were plated (3e3 cells/well) in 96 well plates and treated with a serial dilution of antibody (400, 200, 100, 50, 25, 12.5, 6.25, 3.12, and 1.56nm). Plates were incubated for 84h before cell viability determination was performed with CelTiter-Glo 2.0. Cells exposed to 200nM and 400nM TCRm1-Exatecan had significantly reduced viability (~30% and ~5% respectively) relative to all concentrations of the IgG isotype control-Exatecan tested in this assay. $IC_{50}$ = 194.7nM.

##### *Sequence Pseudo-Perplexity*

For a protein sequence $x_1, \dots, x_N$ we computed sequence pseudo-perplexity (PPPL) as the exponential of the mean per-position token loss,

$$\mathrm{PPPL} = \exp\left(\frac{1}{N}\sum_{i=1}^{N} \ell_i\right),$$

where the per-position loss $\ell_i$ was defined according to one of five protocols. Special tokens (CLS, EOS, padding) were excluded from the mean. For sequences exceeding the model's maximum positional embedding, scores were computed over overlapping sliding windows of length max_length with a stride of max_length / 2, and per-position losses were averaged across overlaps before exponentiation.

Standard masked PPPL[60] defined per-position loss as the negative log probability of the true residue given all other positions masked one at a time: $\ell_i = -log\, p_\theta(x_i \mid x_{\setminus i})$. For efficiency, mask positions were batched diagonally so that each forward pass scored a chunk of consecutive positions simultaneously.

Naive PPPL used the same expression but conditioned on the full unmasked sequence, $\ell_i = -log\, p_\theta(x_i \mid x)$, evaluated in a single forward pass. Because the model observes the target residue in its own input, naive PPPL exhibits substantial information leakage and was retained only as a diagnostic baseline.

Sampled PPPL scored the model on a random 15% subset of positions in a single forward pass, following the position-selection rule of the ESM-2 masked-language-modeling pretraining objective. A subset $S \subseteq \{1, \ldots, N\}$ of valid positions was drawn with selection probability $p_{sel}$ = 0.15. The implementation supports the full pretraining-style replacement scheme in which selected positions are independently masked, replaced with a uniformly random standard amino acid, or left unchanged; for the analyses reported here all selected positions were masked.

Multi-sampled PPPL extended the sampled variant by generating multiple sampled-mask realizations of the same sequence in a single batch (8 realizations in the reported runs). Selected positions were allocated row-wise so that every valid sequence position appeared as a masked position in at least one realization, guaranteeing complete coverage at finite batch sizes. Per-position losses were averaged across the realizations that covered each position before exponentiating the mean.

Direct PPPL replaced the masked inference loop with a trained regression head $f_\phi$ that predicts per-position masked losses from the ESM-2 final-layer hidden states in a single forward pass,

$$\hat{\ell}_i = f_\phi(h_i), \qquad \widehat{\mathrm{PPPL}} = \exp\left(\frac{1}{N}\sum_{i=1}^{N} \hat{\ell}_i\right).$$
